# Exploring genetic interaction manifolds constructed from rich phenotypes

**DOI:** 10.1101/601096

**Authors:** Thomas M. Norman, Max A. Horlbeck, Joseph M. Replogle, Alex Y. Ge, Albert Xu, Marco Jost, Luke A. Gilbert, Jonathan S. Weissman

## Abstract

Synergistic interactions between gene functions drive cellular complexity. However, the combinatorial explosion of possible genetic interactions (GIs) has necessitated the use of scalar interaction readouts (e.g. growth) that conflate diverse outcomes. Here we present an analytical framework for interpreting manifolds constructed from high-dimensional interaction phenotypes. We applied this framework to rich phenotypes obtained by Perturb-seq (single-cell RNA-seq pooled CRISPR screens) profiling of strong GIs mined from a growth-based, gain-of-function GI map. Exploration of this manifold enabled ordering of regulatory pathways, principled classification of GIs (e.g. identifying true suppressors), and mechanistic elucidation of synthetic lethal interactions, including an unexpected synergy between *CBL* and *CNN1* driving erythroid differentiation. Finally, we apply recommender system machine learning to predict interactions, facilitating exploration of vastly larger GI manifolds.

**One Sentence Summary:** Principles and mechanisms of genetic interactions are revealed by rich phenotyping using single-cell RNA sequencing.

## Introduction

A major principle that has emerged from modern genomic and gene expression studies is that the complexity of cell types in multicellular organisms is driven not by a large increase in gene number but instead by the combinatorial expression of a surprisingly small number of components (*1*). This is possible because specific combinations of genes exhibit emergent properties when functioning together, enabling the generation of many diverse cell types and behaviors. Understanding such genetic interactions has important practical and theoretical applications. For example, they can reveal synthetic lethal vulnerabilities in tumors, identify suppressors of inherited and acquired disorders, guide the design of cocktails of genes to drive differentiation between cell types, inform the search for missing inheritance in genetic studies of complex traits, and enable systematic approaches to define gene function in an objective and principled manner (*2*–*5*). Defining how genes interact is thus a central challenge of the post-genomic era.

Given the virtually infinite number of possible combinations and diversity of outcomes, it remains a major challenge to systematically and quantitatively measure combinatorial gene functions in a scalable and information-rich manner. Classically, efforts have focused on measuring pairwise genetic interactions (GI), defined as the extent to which perturbing one gene affects the phenotypic consequences (typically growth rate or viability) of perturbing a second gene (*6*–*10*). Pioneering efforts in yeast have demonstrated that large-scale maps of GIs reveal a unique buffering and synthetic sick/lethal (SSL) GI profile for each gene, which further enables prediction of that gene’s function by clustering of GI profiles (*2*–*4*, *6*, *7*, *11*–*17*). More recently, human GI mapping platforms have enabled unbiased characterization of human gene function (*18*–*28*).

A limitation of most efforts at GI mapping to date is their dependence on univariate readouts of phenotype with low information content (e.g., growth rate), such that GIs are necessarily classified as simply buffering or SSL (i.e., more/less fit than expected). This obscures critical information about how gene combinations lead to emergent properties. For instance, the reprogramming of a pluripotent cell to a terminally differentiated one would modulate the observed growth phenotype as much as altering its metabolism. Put simply, there are many ways for cells to be equally unfit, making it challenging to understand the mechanistic or molecular basis for any particular interaction.

Emerging approaches for rich phenotyping of individual cells present a potential solution to these problems. For example, there is a much more direct relationship between a cell’s “phenotype” in a general sense and its transcriptional state than there is with its growth rate (*5*). We and others have recently developed the Perturb-seq approach, which allows libraries of CRISPR-mediated genetic perturbations to be introduced to cells in pooled format and then measured by single-cell RNA sequencing (*29*–*32*). Each cell is then in effect an independent experiment, enabling highly parallelized studies linking perturbations of individual genes or gene combinations—mediated by single or multiple sgRNAs—to their transcriptional consequences. Rapid advances in single-cell sequencing technologies have enabled Perturb-seq on ∼100,000 cells in a single experiment, and million-cell-scale experiments are already feasible (*29, 33, 34*). This technique thus has the potential to address both the limitations on information content and the scaling challenges of profiling GIs for single and multi-gene perturbations (*29*, *30*).

Here, we present an experimental and computational approach for analyzing and interpreting genetic interactions using the transcriptome as a high-dimensional phenotypic readout. Using CRISPR-mediated activation (CRISPRa) (*35*), we constructed a large-scale, gain-of-function GI map in human cells of genes that modified cell fitness and then broadly sampled the interaction landscape with Perturb-seq. This ensemble of Perturb-seq measurements describes the structure of a manifold—a nonlinear, high-dimensional surface—populated by transcriptional phenotypes. The manifold, analogous to Waddington’s canal in differentiation, represents the cell states reachable from a given starting point through genetic perturbations. The local shape of the manifold thus reflects how genes interact, so we refer to it as a GI manifold. This high-dimensional, geometric approach to GIs allows one to generalize the concepts of buffering and SSL interactions to reveal critical functional distinctions previously obscured by describing GIs in a single dimension.

Specifically, we found the use of rich phenotypes enabled us to distinguish cell death, slow growth, and differentiation to a variety of cell states as mechanisms underlying fitness-level GIs. Exploration of the GI manifold also allowed us to determine the order of genes in linear pathways and identify distinct modes of interaction—for example, by distinguishing between epistatic relationships and genetic suppressors that could each underlie buffering interactions. Finally, we applied machine learning on the transcriptional phenotypes to predict fitness-level interactions, an approach we propose to further mitigate the scaling challenges of screening for novel GIs in a sparse landscape. We expect the conceptual and computational frameworks presented here will be broadly applicable to genetic interactions obtained via other rich phenotyping approaches (e.g. proteomics, imaging) and methods of perturbation (e.g. knockdown, knockout, mutagenesis, chemical perturbations).

## Results

### A CRISPRa fitness-level genetic interaction map

In order to generate a large and diverse set of gene pairs for examination by rich phenotyping, we focused on genes that modified cell fitness when overexpressed. We chose fitness since it is a complex and readily measured phenotype that integrates many different physiological processes. We chose gain-of-function perturbations as they are an effective way of exploring cell states (*5*, *36*–*38*), and with CRISPRa are now feasible to implement systematically for quantitative interaction studies. From a previous genome-wide CRISPRa screen, we selected 112 hit genes whose activation enhanced or retarded growth of K562 cells (Fig. 1A) (*35*). As strong GIs are expected to be sparse in the overall GI landscape (*2*, *3*), we reasoned that pairs of genes with primary effects were more likely to strongly interact when combined.

**Figure 1.**
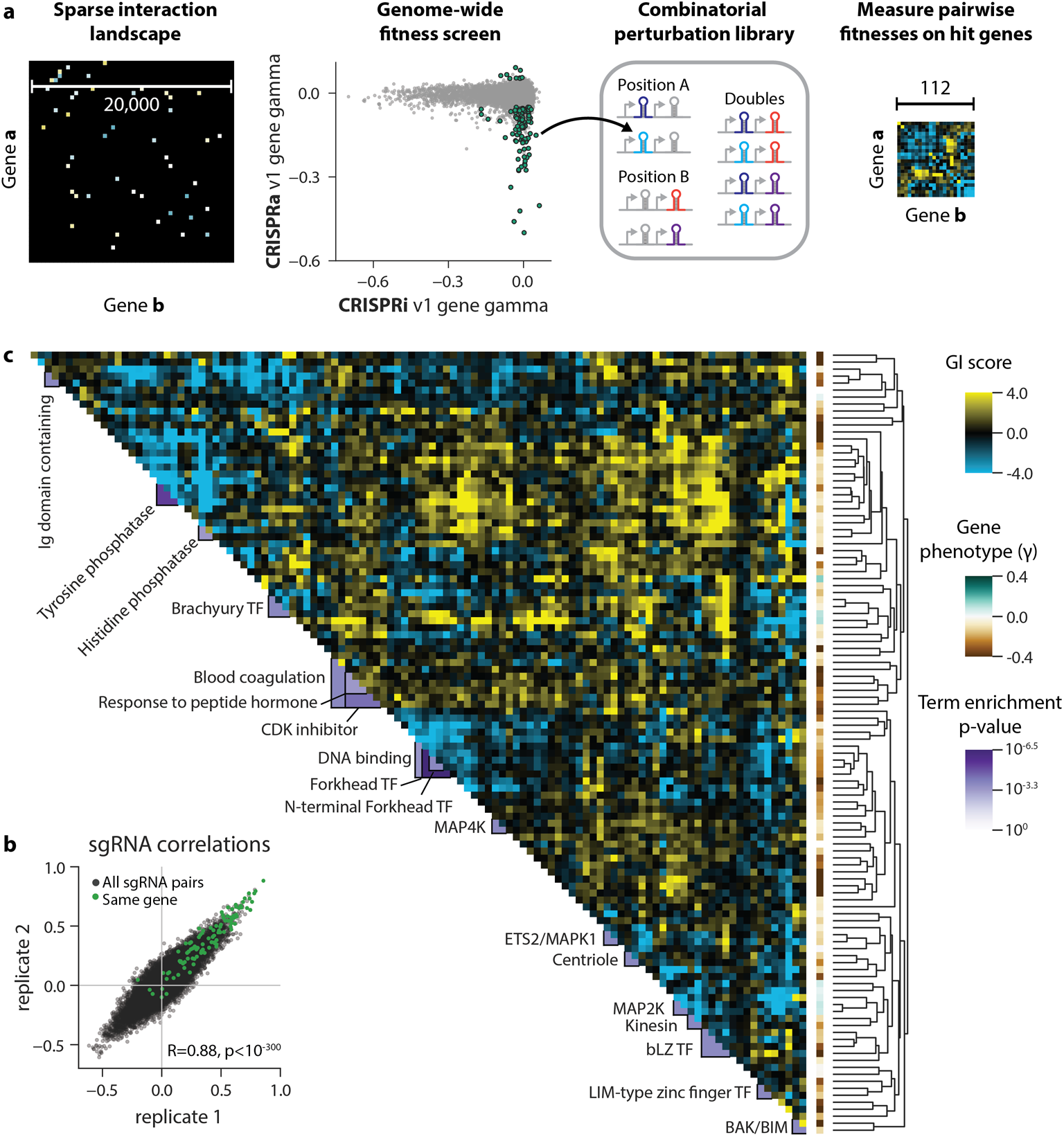
A CRISPRa fitness-level genetic interaction map. **(A)** Construction of a CRISPRa GI map. (Left) A theoretical genome-wide genetic interaction map is sparsely populated with strong genetic interactions. (Middle) Genes with a fitness phenotype upon overexpression with CRISPRa were selected from a genome-scale screen performed in K562, the majority of which did not have a fitness phenotype upon repression with CRISPRi (*35*). Two sgRNAs targeting each gene were cloned into a lentiviral dual sgRNA expression library. (Right) A GI map was constructed by quantifying sgRNA pair fitness phenotypes in K562 relative to the expected pair phenotypes. **(B)** sgRNA GI correlations from two independent replicates. sgRNAs targeting the same gene had high GI profile correlations relative to all sgRNA pairs (see also Figure S1D). **(C)** CRISPRa fitness-level GI map. Gene-level GIs were clustered by average linkage hierarchical clustering based on Pearson correlation between genes. Clusters were annotated by assigning DAVID annotations if a DAVID term was significantly enriched in that cluster (hypergeometric ln(p) ≤ −7.5) and more enriched than in any other cluster (see Methods).

To systematically measure these interactions, we adapted a technology we previously developed for constructing genetic interaction maps in human cells using CRISPRi (*18*) (fig. S1A-B). Briefly, each candidate interaction was probed by constructing a cassette containing two sgRNAs targeting the two constituent genes along with a unique barcode sequence. Each gene was targeted using two distinct sgRNAs, generating a genetic interaction (GI) library composed of 57,121 cassettes in total (28,680 unique sgRNA pairs). K562 cells stably expressing the SunTag CRISPRa system (*39*) were transduced with the CRISPRa GI library by pooled lentiviral transduction, and sgRNA pair abundance was compared at the start of the screen and after ten days of growth to infer fitness phenotypes (expressed as γ, the log_2_ enrichment of sgRNA pair abundance scaled by number of cell doublings in the screen). We used a custom sequencing and analysis pipeline that corrects for lentiviral recombination events ((*18*); see methods; fig. S2, A and B) and calculates sgRNA GIs by measuring deviation relative to the average trend determined by a quadratic fit of fitness measurements (fig. S2C). Gene-level GIs were computed by averaging all sgRNA pairs targeting a given interaction. We found high sgRNA- and gene-level GI concordance between independent replicates (gene-level GI R=0.80, p<10^−300^; Fig. S2D). Importantly, the GI profiles for sgRNAs targeting the same gene were much more similar than the background of all sgRNA GI correlations (median R=0.50 compared to 0.04; Fig. 1B and fig. S2, D, E and F).

We then clustered genes according to the similarity of their GI profiles to produce a GI map (Fig. 1C; larger version with gene labels provided in fig. S3A). The map had low-rank structure: that is, groups of genes interact similarly so that there are fewer overall degrees of freedom than genes, resulting in block-like structure. Unbiased DAVID term annotation showed that multiple blocks corresponded to known biological units of organization (*40*, *41*), such as forkhead box-containing transcription factors, cell cycle inhibitors, and the direct physical interactors *PLK4* and *STIL* (*42*). The map also shed light on some genes of unknown function. For example, the transcription factor *ZBTB10* clustered alongside Snail (*SNAI1*) and *DLX2*, suggesting a possible role in epithelial-to-mesenchymal transition (*43*). We therefore conclude that CRISPRa GI maps, like past efforts based on loss-of-function alleles (*18*), can assign function to individual genes by the similarity of GI profiles.

Additionally, we found that individual GIs in the CRISPRa GI map can generate hypotheses consistent with known biology. For instance, the pro-apoptotic factors Bak and Bim (*BAK1* and *BCL2L11*) are strongly buffering (Fig. 1C), as they function in a linear signaling pathway to promote cell death and upregulation of either factor is sufficient to elicit a strong growth defect (*44*). However, the origins of many specific interactions were difficult to deduce, as each interaction is characterized only by a single scalar value in a fitness-level map. Furthermore, not all the large-scale hierarchical structure of the clustered map could be readily explained. Therefore, although GI mapping allows us to efficiently screen many candidate interactions to identify those that are surprising, higher resolution is needed to understand the biological underpinnings of specific interactions and correlations.

### Perturb-seq reveals diverse phenotypes driving growth defects

Ultimately, the limitation with both loss- and gain-of-function GI maps is that one-dimensional readouts such as fitness are a coarse readout of a cell’s state. If cellular phenotype in a broad sense is thought of as lying on a high-dimensional manifold, different genetic perturbations may push that cell to drastically different positions that nevertheless result in similar fitness defects (Fig. 2A). Rich phenotyping techniques such as Perturb-seq present a potential solution to this problem by providing a more complex measure of cellular phenotype, allowing us to define the structure of this manifold (*29*).

**Figure 2.**
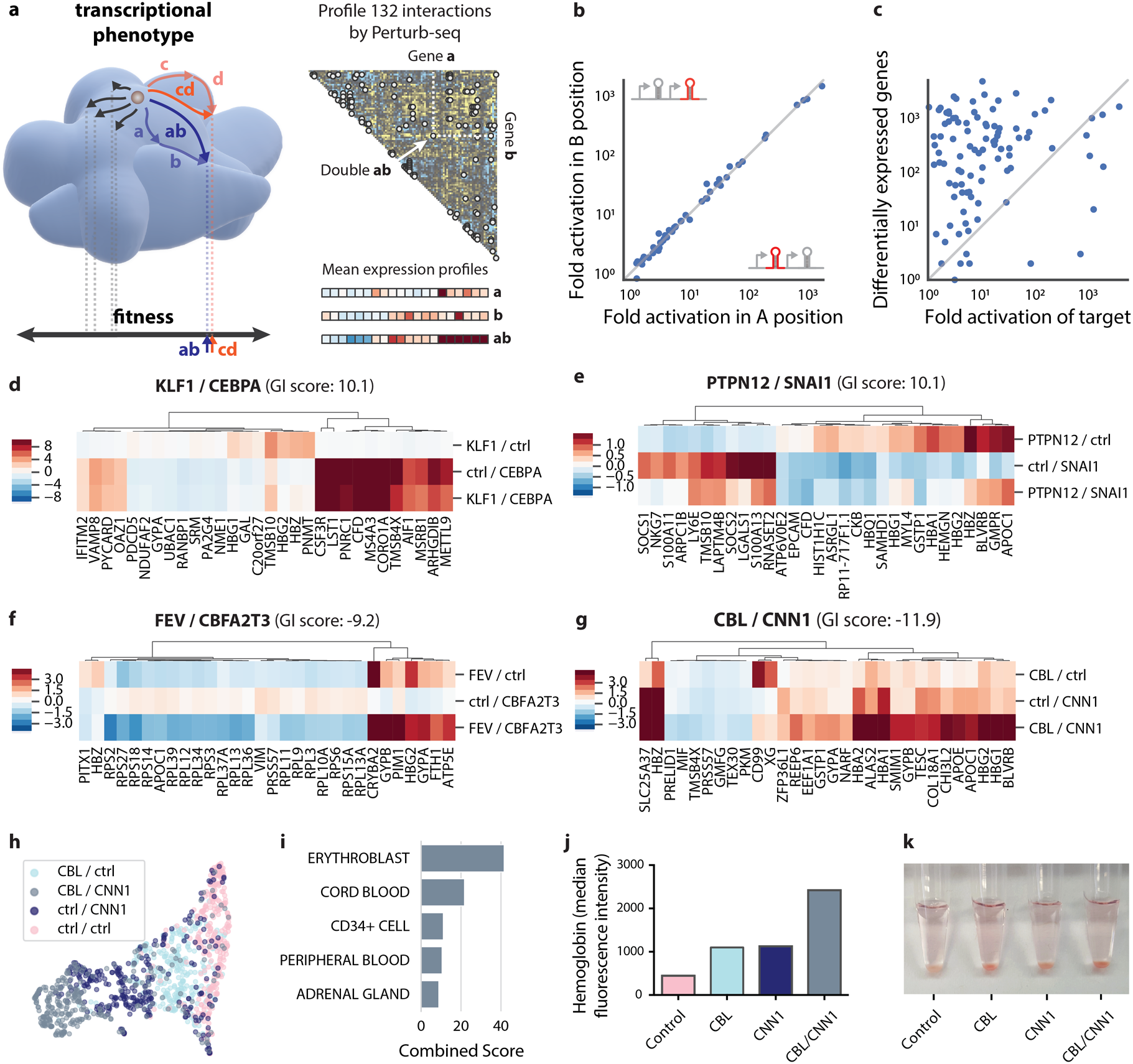
Dissecting genetic interactions using rich phenotypes. **(A)** Representation of the GI manifold model. (Left) A given cell’s transcriptional state defines a point on a high-dimensional surface. Genetic perturbations push cells to different points on this manifold. Distant points on the manifold may result in similar scalar fitness measurements. (Right) Sampling of pairs from GI map for profiling by Perturb-seq. **(B)** Comparison of fold activation of target genes when the targeting sgRNA is in the A or B position in the dual sgRNA expression cassette. **(C)** Fold activation of the target gene compared with the total number of differentially expressed genes. **(D)-(G)** Examples of rich phenotyping of GIs with Perturb-seq. For each GI, average transcriptional profiles for the two constituent single perturbations are compared to the double perturbation. Heatmaps show deviation in gene expression relative to unperturbed cells. **(H)** UMAP projection of single-cell Perturb-seq data in the *CBL*/*CNN1* interaction. Each dot is a cell colored according to genetic background. **(I)** ARCHS4 cell type term enrichment for genes showing large expression changes in CBL/CNN1 doubly-perturbed cells (*66*). **(J)** Expression of hemoglobin in HUDEP2 cells upon cDNA overexpression of *CBL* or *CNN1*. Hemoglobin was labeled with anti-HbF antibody and measured by flow cytometry. **(K)** Pelleted HUDEP2 cells. Hemoglobin expression appears red.

We thus set out to profile genetic interactions via Perturb-seq. Given the low-rank structure of the GI map, we reasoned that we could sample most of the biology present in our GI map without measuring all gene pairs, as many blocks of interactions are likely explained by similar mechanisms. We picked 132 gene pairs from throughout the map, chosen both within and between blocks of genes with similar interaction profiles, and targeted each with CRISPRa sgRNA pairs (Fig. 1A and fig. S4A). We also profiled all single gene perturbations to enable direct comparison of individual perturbations and genetic interactions (i.e. single gene A, single gene B, and pair AB). In total we obtained transcriptional readouts for 287 perturbations measured across ∼110,000 single cells (Fig. 2A, fig S1C-D, and Table S7).

Using these data, we first assessed the performance of our CRISPRa reagents (Table S7). The levels of target gene activation spanned a broad range both in terms of fold-change from baseline expression level (Fig. 2B and fig. S4B) and of absolute transcript abundance (fig. S4C). As a general trend, genes that were poorly expressed could be induced more than those that were highly expressed, similar to previous findings (Fig. S4, B and C) (*45*), although this did not explain most of the variation in activation. Expression of genes neighboring the target was generally unperturbed with the exception of transcripts that shared promoter regions (Methods; fig. S5, Table S8). The A and B positions of the sgRNA cassette produced similar activation levels (Fig. 2B), number of differentially expressed genes (Fig. S4D), and highly-correlated transcriptional profiles (Fig. S4F). Finally, there was no correlation between fold activation and the number of differentially expressed genes (R=0.07; Fig. 2C), implying that even a relatively small increase in the mRNA abundance of a target gene can radically alter a cell’s state. We did observe a relationship between the number of differentially expressed genes and the degree of fitness defect (fig. S4E).

As with GI profiles, Perturb-seq profiles of single gene perturbations can be used to cluster genes and predict functional associations (*29*). To compare this gain-of-function Perturb-seq map, we applied the same GI clustering and annotation pipeline to all pairwise Pearson correlation coefficients between single gene overexpression profiles (fig. S6A). This map largely recapitulated many of the DAVID-annotated clusters identified in the GI map. Although there was overall concordance between GI and Perturb-seq correlations (R=0.29, p<10^-103^; fig. S6B), for some genes the transcriptional phenotype produced a far broader range of correlations (e.g. *CDKN1A*) and vice versa (e.g. *BAK1*; fig. S6, C and D), indicating that transcriptional and fitness maps produce complementary information.

To explore the ability of rich phenotyping to better resolve genetic interactions, we examined examples of buffering and SSL interactions that exhibited similar GI scores in the fitness-level map but appeared to behave differently on a transcriptional level. We first considered two strong buffering interactions that each had GI scores of +10.1, *KLF1*/*CEBPA* and *PTPN12*/*SNAI1*. (GI scores were approximately normally distributed and ranged from −18.0 to +11.5 with 5^th^ and 95^th^ percentiles −3.5 and +3.3; Fig. S3B.) The buffering GI between *KLF1*/*CEBPA* is explained by near complete genetic epistasis of *CEBPA* over *KLF1* (Fig. 2D). Meanwhile, the interaction between *PTPN12*/*SNAI1* (Fig. 2E) results from genetic suppression, as the GI transcriptional signature is a superposition of weaker versions of the two single gene perturbations. We observed similarly diverse behaviors among SSL interactions. For example, in the *FEV*/*CBFA2T3* interaction (GI score: −9.2), the addition of the otherwise weak *CBFA2T3* perturbation appeared to strongly potentiate the effects of the *FEV* perturbation (Fig. 2F). Meanwhile, combining perturbations of *CBL* and *CNN1* (calponin) (GI score: −11.9) resulted in a profile that resembled but exceeded the additive expectation (Fig. 2G and fig. S7A). These results demonstrate that rich phenotyping identifies modes of interaction that are missed by scalar GI measurements.

We were intrigued by the unexpected SSL interaction between *CBL* and *CNN1*. *CBL* is a negative regulator of receptor tyrosine kinase signaling (*46*). *CNN1* is a poorly characterized gene that is annotated as a smooth-muscle-specific protein, although *CNN1* is expressed in many cell types (*47*–*49*). Single-cell analysis revealed an apparent progression of phenotype from unperturbed through to doubly-perturbed *CBL*/*CNN1* cells (Fig. 2H). Among the most strongly induced genes in the doubly-perturbed cells were hemoglobin genes (6–39-fold), an iron importer involved in heme biosynthesis (*SLC25A37*, 13-fold), and the blood group antigen CD235a (*GYPA*, 2-fold) (Fig. 2G). Enrichment analysis of these markers and others showed similarity to an erythroblast transcriptional signature (Fig. 2I), suggesting that rather than arising from cell death, this interaction is driven at least in part by erythroid differentiation and a concomitant decrease in proliferation. Consistent with this finding, K562 cells have been widely used to study erythroid and other myeloid differentiation programs (*50*, *51*).

To validate this hypothesis, we turned to cDNA overexpression experiments in K562 cells and in HUDEP2 cells, an immortalized human erythroid progenitor cell line that can undergo erythroid differentiation and enucleation (*52*). We transduced each cell line with lentiviral vectors encoding *CBL* and *CNN1* cDNAs alone and in combination, and again observed strong induction of markers of erythroid differentiation (Fig 2J and fig. S7, B and C). Even under HUDEP2 self-renewal conditions that normally do not support erythroid differentiation, these cells produced hemoglobin to the extent that centrifuged cell pellets turned red, suggesting this induced differentiation program dominates over self-renewal signaling (Fig. 2K). Together, these results show how Perturb-seq analysis can quickly lead to a hypothesis about the biology underlying a genetic interaction even when one of the components is poorly understood.

### Constructing a high-dimensional GI manifold

Encouraged by our examples showing that Perturb-seq could identify both how perturbations combined and the mechanisms underlying specific genetic interactions, we next turned to a global view. Geometrically, each possible cellular transcriptional state defines a point on the surface of the GI manifold. By applying and analyzing a sufficiently diverse set of perturbations, the shape of this surface can then be inferred. Given the low-rank nature of our GI map, we reasoned that our set of perturbations sampled many of the states reachable from a control population, enabling us to construct a representative view of the GI manifold governing fitness-level GIs in K562 cells (Fig. 3A).

**Figure 3.**
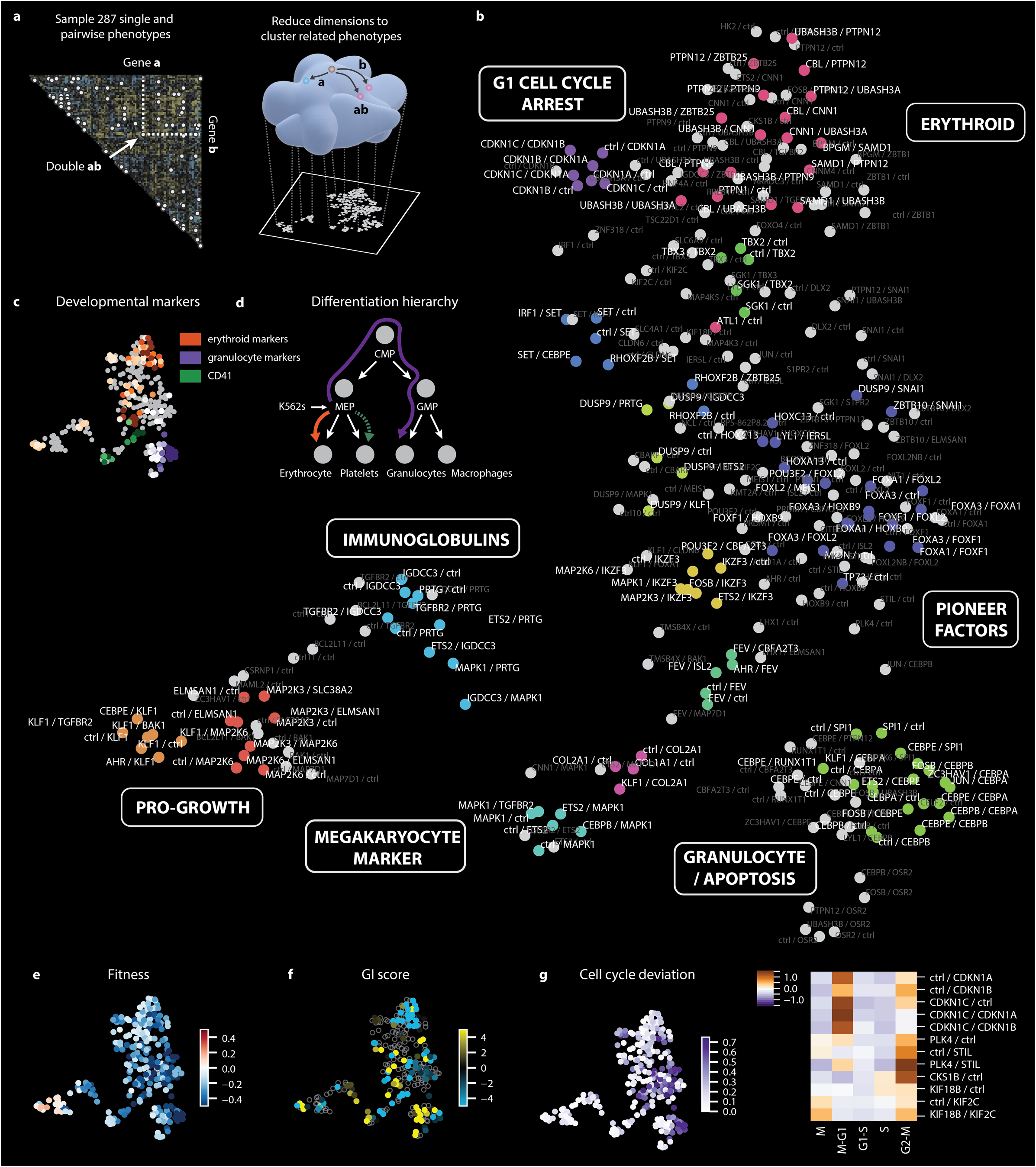
Visualization of the GI manifold. **(A)** Using diverse genetic perturbations, the structure of the GI manifold can be inferred and then visualized by dimensionality reduction to a plane. **(B)** UMAP projection of all single gene and gene pair Perturb-seq profiles. Each dot represents a genetic perturbation. Clusters of transcriptionally similar perturbations are colored identically, while grey dots are genes that do not fall within stable clusters. **(C)** Expression of marker genes for different hematopoietic cell types. Color is scaled by mean expression Z-score of the marker gene panel. **(D)** Hematopoietic differentiation hierarchy. K562 cells are a poorly differentiated erythroid cell line. **(E)** Fitness measurements from the GI map, expressed as gene pair growth phenotypes (γ). **(F)** GI scores from the fitness-level GI map. Single gene perturbations are not included. **(G)** Cell cycle phenotypes. (Left) Cell cycle deviation scores. Stronger scores indicate alteration from the distribution of cell cycle positions observed in unperturbed cells. (Right) Relative enrichment or depletion of cell cycle phases relative to unperturbed cells induced by selected genetic perturbations.

To visualize this GI manifold, we used UMAP (uniform manifold approximation and projection) to project our 287 mean expression profiles into a two-dimensional plane (Fig. 3B) (*53*). This projection identified groups of genetic perturbations driven by similar mechanisms. For example, the *CBL*/*CNN1* SSL interaction was one of many inducing erythroid differentiation, thus explaining a large block of activity in the GI map (Fig. 3, B to D and fig. S8). Most of these interactions surrounded *CBL*, its regulators *UBASH3A*/*B*, and several multi-substrate tyrosine phosphatases (e.g. *PTPN9*/*12*). Given the role of these genes in regulating receptor tyrosine kinases, one possible explanation for their overall behavior is that reduction in BCR-ABL signaling, which is the driver oncogene in K562 cells, induces an erythroid differentiation program (*50*, *51*).

We inspected our data to look for evidence of perturbations promoting other differentiation states. Additional groups of perturbations were characterized by their expression of granulocyte and megakaryocyte markers (Fig. 3C and fig. S8), suggesting that our perturbations explore multi-lineage potential (Fig. 3D). For example, the granulocyte cluster mostly involved canonical regulators such as C/EBP-α, -β, -ε (*CEBPA/B/G*) and PU.1 (*SPI1*) (*54*). These perturbations also significantly lowered fitness and increased production of both pro-and anti-apoptotic genes (fig. S8G). Intriguingly, the cluster expressing the canonical megakaryocyte marker CD41 did not appear transcriptionally similar to differentiating megakaryocytes, and did not adopt any of the characteristic morphological features of megakaryocytes by microscopy (fig. S8H and I), suggesting that these cells are perhaps only primed towards megakaryocytic differentiation (*55*, *56*). This result highlights how dependence on single marker gene expression to define a cell’s state can be misleading (*57*). Nevertheless, primed cells may still be a useful starting point for future screens with additional perturbations. By simultaneously detecting signatures of multiple differentiation states, our approach can support and enhance higher-order combinatorial perturbation screens aimed at improving protocols for driving cells into distinct differentiation states.

We also observed two clusters driven by perturbations that increased fitness, arising either from interactions with *MAP2K3*/*6* (and a likely role in p38 MAPK signaling) or the major erythroid developmental regulator *KLF1* (Fig. 3E). More generally, however, both fitness and GI scores (Fig. 3E-F) were distributed throughout the GI manifold, consistent with the idea that univariate fitness measurements collapse the much larger landscape of transcriptional states.

The projection of the GI manifold is ultimately the product of ∼110,000 single-cell transcriptomes, and so it is also sensitive to single-cell phenotypes. For example, we observed that a group of interactions involving CDK inhibitor genes all caused arrest in G1, *CKS1B* overexpression and the *PLK4*/*STIL* interaction induced arrest in G2/M, and the *KIF18B*/*KIF2C* interaction (a previously identified physical interaction for which we measured a GI score of − 10.7) caused accumulation in M phase (Fig. 3G) (*58*). These phenotypes are all consistent with the known functions of these genes and demonstrate how our approach is sensitive to both first-(mean) and second-(variance) order properties of phenotypes.

### Quantitative modeling of high dimensional genetic interactions

If each transcriptional state defines a point on the surface of the GI manifold, then each genetic perturbation defines a possible direction in which cells can be pushed (Fig. 4A). Understanding how genes interact therefore corresponds to understanding how the doubly-perturbed state can be reached from the unperturbed state by traveling along some combination of the directions given by the two single perturbations. This can be estimated to first order by fitting a regression model

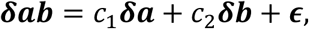

**Figure 4.**
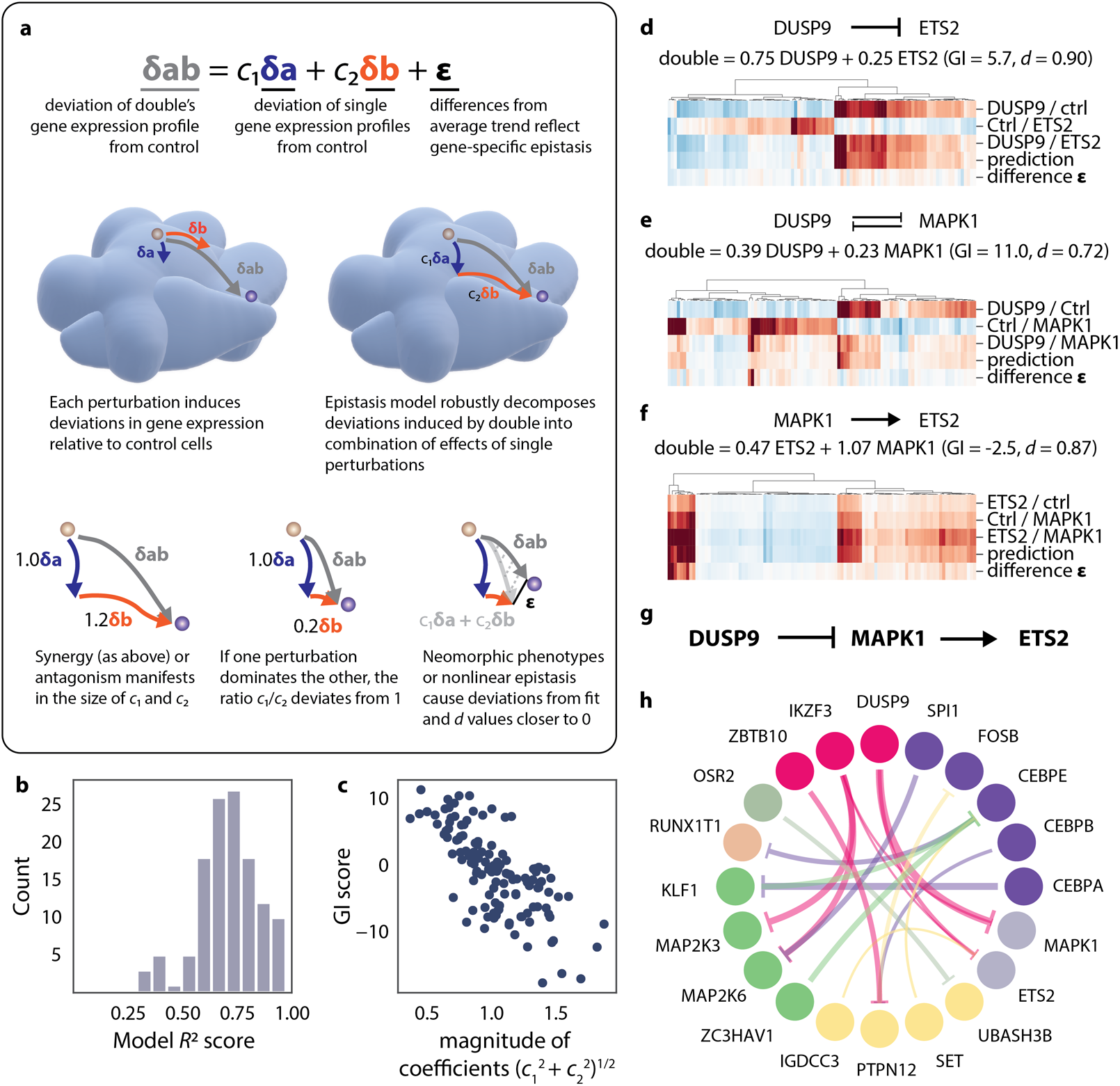
A quantitative model for high-dimensional GIs. **(A)** Model of transcriptional genetic interactions. Different transcriptional states define points on the surface of the GI manifold and genetic perturbations define vectors of travel. The model decomposes double perturbations as a linear combination of the two constituent single perturbations. **(B)** Model fit across all measured double perturbation Perturb-seq profiles. **(C)** Magnitude of model coefficients in Perturb-seq experiment compared to GI score from the fitness-level GI map. **(D)-(F)** Application of the model to selected GIs measured using Perturb-seq. For each GI, transcriptional profiles for the two constituent single perturbations are compared to the double perturbation, the model fit, and the model residual. Heatmaps show deviation in gene expression relative to unperturbed cells. **(G)** Model of genetic interactions among *DUSP9*, *MAPK1*, and *ETS2* inferred from model fits. **(H)** Epistatic buffering interactions oriented using the genetic interaction model. Each arrow denotes a genetic interaction, originating in the gene that dominates when the two genes are simultaneously perturbed. Arrow size denotes the degree of dominance as measured by the asymmetry of model coefficients. Genetic perturbations with similar transcriptional profiles are colored identically.

that represents the transcriptional effect of the combined perturbation ***δab*** in terms of the two single perturbations (Table S9). To minimize the influence of outlier genes, we fit this model by robust regression (Methods). The relative sizes of the coefficients *c*_1_ and *c*_2_ then measure how much of the pair phenotype is accounted for by each single perturbation (Fig. 4A), and the model’s fit, which we quantified via distance correlation (*d*, Methods) measures the amount of novel behavior that arises when combining perturbations.

This simple linear fit model explained more than 70% of the variance in gene expression on average (Fig. 4B and fig. S9A; mean *R*^2^ = 0.71)—markedly better than an additive model of interaction (fig. S9B; mean *R*^2^ = 0.43). As with fitness-based GI scores (*2*, *11*, *18*), overall correlation of expression profiles was somewhat predictive of interactions (*R* = 0.42; fig. S9C).

Notably, we observed a robust anti-correlation (*R* = −0.72) between the magnitude of these coefficients and the fitness-based GI score (Fig. 4C), with larger coefficients indicating SSL interactions and smaller ones buffering interactions. The strength of this relationship was somewhat surprising given that the two measures of interaction were derived by unrelated analysis methods from different datasets. Comparing the two demonstrated that a GI score of 0 corresponded to a coefficient magnitude of 1 (Fig. 4C). An intuitive explanation of the relationship further emphasizes the utility of the GI manifold—buffering interactions travel less “far” than expected both in high-dimensional space and when projected to one-dimensional fitness space, and vice versa for SSL interactions (Fig. 4A).

The model allowed us to quantify how transcriptional profiles combined to yield interactions. For example, buffering interactions could arise either from epistasis, in which one perturbation largely canceled the effects of the other (e.g. *KLF1*/*CEBPA*, *c*_1_ = 0.19, *c*_2_ = 0.72, Fig. 2D and fig. S9D), or via suppression, when two perturbations attenuated each other’s activities (e.g. *PTPN12*/*SNAI1*, *c*_1_ = 0.60, *c*_2_ = 0.57, Fig. 2E and fig. S9E). See also our analysis of *DUSP9*/*MAPK1*/*ETS2* in Figure 4 and 6. By contrast, many synthetic lethal interactions arose from synergy between transcriptionally similar perturbations (e.g. *CBL*/*CNN1*, *c*_1_ = 1.24, *c*_2_ = 0.8, Fig. 2G and fig. S9G). Novel behavior (i.e. lower values of *d*) arose either via potentiation, in which a perturbation that had little transcriptional effect on its own appeared to enhance the effects of a partner (e.g. *FEV*/*CBFA2T3*, *c*_1_ = 1.46, *c*_2_ = 1.17, *d* = 0.74, Fig. 2F and fig. S9F), or by relatively rare instances where double phenotypes were predominantly neomorphic (e.g. *PLK4*/*STIL*, *c*_1_ = 0.42, *c*_2_ = 0.75, *d* = 0.53, fig. S9H).

The model’s parameters (Table S9) thus provide a simple, useful summary of how perturbations combine, essentially acting as a generalization of the one-dimensional “buffering vs. synthetic lethal” paradigm that has typically been used to categorize genetic interactions. However, because the model is derived from rich phenotypes, it provides insights that are obscured with scalar measures of interaction.

One of the simplest benefits that results is that buffering interactions can be oriented based on which single perturbation better explains the double. For example, the *DUSP9* phenotype strongly dominated the *DUSP9*/*ETS2* interaction (GI score: 5.7, *c*_1_ = 0.75, *c*_2_ = 0.25, Fig. 4D), suggesting that DUSP9 directly or indirectly inhibits ETS2 activity. Meanwhile, DUSP9 and MAPK1 clearly antagonized each other’s activities (GI score: 11.0, *c*_1_ = 0.39, *c*_2_ = 0.23, Fig. 4E). Finally, examining the *ETS2*/*MAPK1* interaction (GI score: −2.5, *c*_1_ = 0.47, *c*_2_ = 1.07, Fig 4F) showed that the two perturbations induced similar overall phenotypes. More specifically, the *ETS2* transcript was as overexpressed in *MAPK1*-perturbed cells (9.3 fold) as in *ETS2*-perturbed cells (9.2 fold), and that the two perturbations synergized (35.8 fold overexpression in *ETS2*/*MAPK1*). This behavior, in which a second perturbation acts at least partly by upregulating its interaction partner, was uncommon in our dataset (fig. S9I). Taken together, these three results suggest a linear regulatory pathway in which DUSP9 inhibits MAPK1, which activates ETS2—an example of classical gene epistasis. Our data are consistent with the known roles of these gene families: dual specificity phosphatases such as DUSP9 can dephosphorylate ERK family kinases, such as MAPK1 (Fig. 4G) (*59*). Similarly, ERK family kinases can phosphorylate and activate transcription factors such as the ETS proteins. We further discuss the mutual suppression of *DUSP9* and *MAPK1* in Figure 6 at single-cell resolution. More generally, these results show how rich phenotyping can directly generate hypotheses about gene regulation. Following this logic, we can for example orient all the buffering interactions in which one perturbation is epistatic to another (Fig. 4H).

### Dimensionality reduction reveals structure in genetic interactions

Because the model coefficients are not dependent on specific biological phenotypes, they also provide a means to look for large-scale structure in the types of interactions that exist. We used a two-dimensional visualization technique (*60*) to project properties of each interaction along two axes (Fig. 5A and Table S9; Methods). The first axis was defined by properties of the model: the magnitude of the coefficients (measuring interaction strength), the asymmetry of the coefficients, and the model fit. In the second axis we compared the transcriptional profiles directly: how similar the single profiles were to the double, to each other, and whether one single profile was more similar to the double than the other. The resulting plot (Fig. 5B) showed that the model can act as a type of dimensionality reduction, projecting each interaction onto an archetype of behavior. This analysis demonstrates that the examples previously presented of epistasis (Fig. 2D and 4D and fig. S9D), suppression (Fig. 2E and 4E and fig. S9E), redundancy (Fig. 3G and 4F) and neomorphism (fig. S9H) are representative of structural features of how genes interact within our dataset, and that distinct behaviors can underlie “buffering” and “synthetic” interactions. Since this approach does not depend on specific features of gene expression data, it can also be applied to other high-dimensional readouts, yielding a simple strategy for defining, measuring, and qualitatively characterizing genetic interactions measured with any rich phenotyping technology.

**Figure 5.**
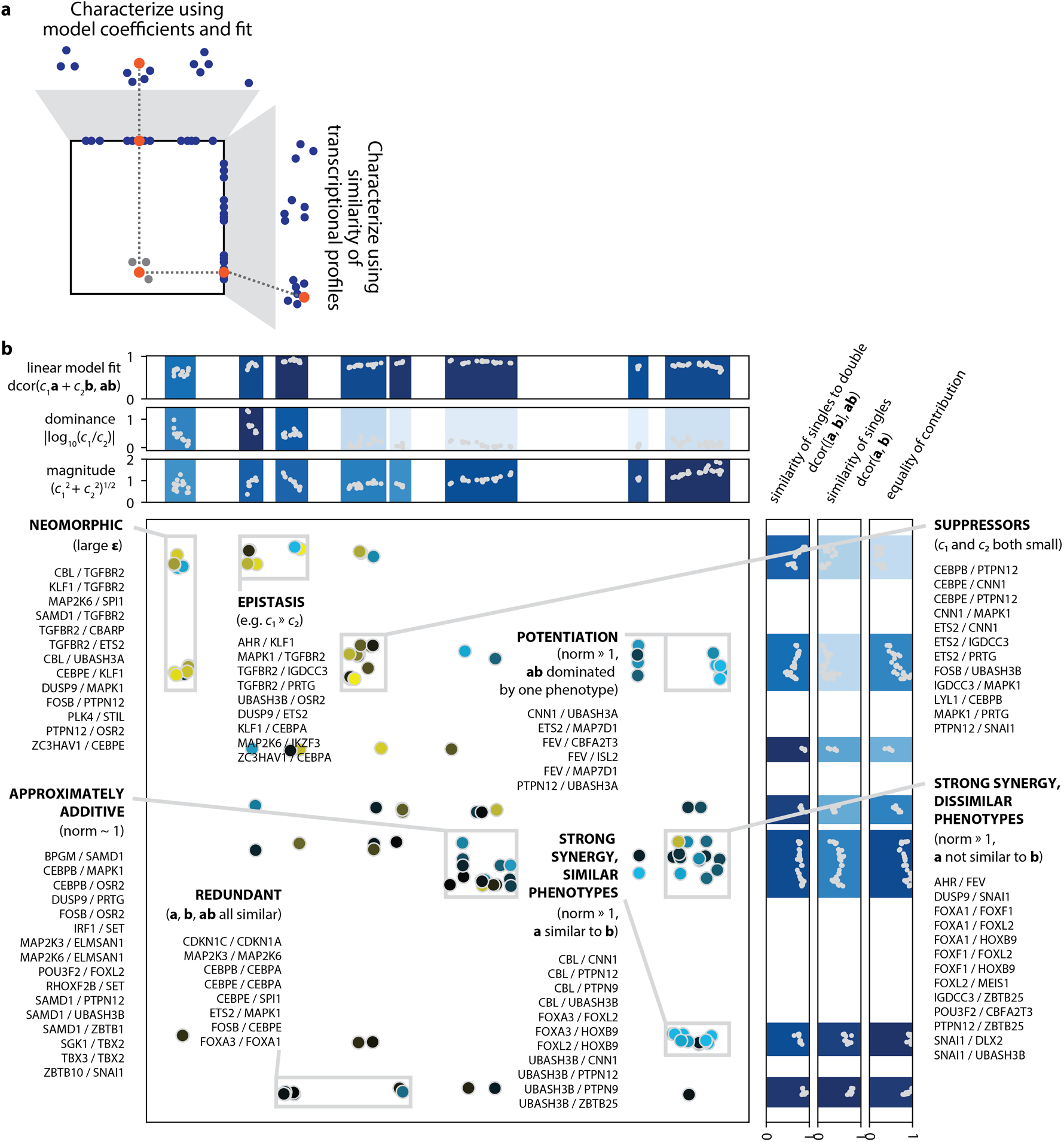
Categorizing GIs using features derived from rich phenotyping. **(A)** Schematic of approach to summarize GIs. Each GI was characterized using features derived from the model (x-axis) and by measures of similarity among the transcriptional profiles (y-axis). These two viewpoints were each clustered and collapsed using UMAP to a single dimension to define the two axes. **(B)** Visualization of all measured GIs in Perturb-seq experiment. The features defining the two axes are plotted next to them. Categories of GIs are annotated based on features shared within the clusters.

### Single-cell heterogeneity reveals the trajectory of genetic interactions

The assertions made by our model above are derived from an averaged phenotype for each perturbation. We reasoned that in some instances, exploiting the single-cell resolution of our measurements would provide more information, as heterogeneity among cells with the same perturbation would allow a wider swath of phenotypic space to be measured (Fig. 6A).

**Figure 6.**
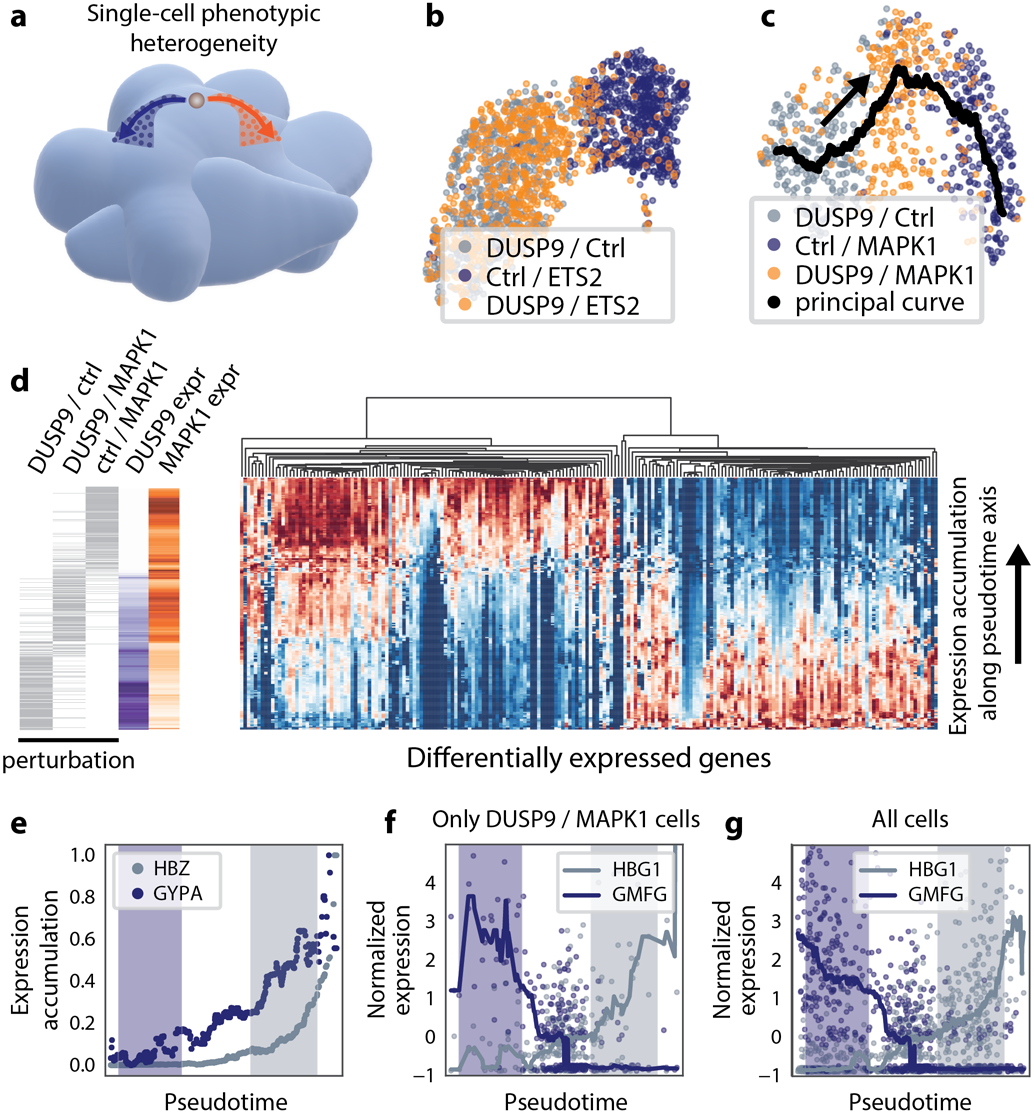
Single-cell trajectories of GIs. **(A)** Stochastic heterogeneity can cause individual cells (dots) bearing a given genetic perturbation to explore the space on the GI manifold surrounding the average direction of travel (arrows). **(B)** UMAP projection of single cells with overexpression of *DUSP9* and/or *ETS2*. Cells with *DUSP9* overexpression overlap with *DUSP9*/*ETS2*, largely recapitulating the population average behavior (Fig. 4D). **(C)** UMAP projection of single cells with overexpression of *DUSP9* and/or *MAPK1*. Black line represents the principal curve, which tracks the primary direction of variation in the dataset and defines a scalar “pseudotime” that can be used to order all cells. **(D)** Gene expression averaged along the principal curve. Each row denotes a cell ordered according to position along the principal curve. The left three columns indicate that cell’s genetic background. At each point, cells that are close on the principal curve are averaged to produce a local estimate of median gene expression. The heatmap shows normalized expression of differentially expressed genes. The *DUSP9* and *MAPK1* expression columns show the same data for the targeted genes. **(E)** Accumulation profiles of two highly expressed marker genes with different behaviors. **(F)** Expression profiles of two highly expressed markers averaged over those cells in (C) bearing the DUSP9/MAPK1 double perturbation **(G)** The same profiles averaged over all cells in (C).

For example, further inspection of the *DUSP9*/*MAPK1*/*ETS2* interactions showed that the single-cell data largely recapitulated the average behavior for the *DUSP9*/*ETS2* (Fig. 6B) and *MAPK1*/*ETS2* (fig. S10A) interactions. However, cells bearing *DUSP9*/*MAPK1* perturbations showed a range of phenotypes in which doubly-perturbed cells comingled with and bridged the gap between cells bearing the two single perturbations (Fig. 6C). This behavior may reflect direct antagonism of *MAPK1* by *DUSP9*, so that small differences in levels of activation lead to relatively large differences downstream because each additional active kinase molecule has a relatively large effect.

To investigate this antagonism, we defined a pseudotime axis ((*61*); Methods) along the direction of principal variation and averaged cell expression profiles along this. Examining differentially expressed genes (Fig. 6D) showed an apparent continuum of phenotype. Highly expressed marker genes, which can be more reliably measured in each individual cell, showed distinctly different patterns of accumulation along this axis (Fig. 6E). Consistent with the UMAP projection, these markers showed similar expression patterns within only doubly-perturbed cells (Fig. 6F) as they did within all cells considered (Fig. 6G), suggesting that phenotypic heterogeneity within doubly-perturbed cells led to individual cells with phenotypes that were indistinguishable from cells bearing each of the two single perturbations. This variation did not appear to be the result of stable differences in the expression levels of *MAPK1* and *DUSP9* (fig. S10B). *DUSP9* and *MAPK1* therefore appear to oppose each other’s activity entirely and thus are true suppressors. As each of these enzymes can be regulated by many factors (*59*), a scalable assay that can be used to determine which interactions are strongest in a given context is a useful means to understand cell type-specific kinase and phosphatase dependencies.

### Predicting pairs of interacting genes using a recommender system

Regardless of experimental approach, any study of GIs must ultimately confront the challenge posed by their intractable scaling in the number of genes considered. Because of the low-rank structure of the GI map, we hypothesized that an optimal route to explore GI space could be to measure a subset of interactions at a fitness level and computationally predict the remainder.

We reasoned that there is a substantial similarity between this problem and that of predicting a person’s shopping preferences based on past buying behavior, a so-called “recommender system.” Recommendations are typically made in a regime in which measurements are very sparse, as each user has generally purchased only a tiny fraction of items (*62*). A critical component in these approaches is that they can leverage side information obtained by other means to improve predictive power. To apply this framework to genetic interactions, we used the similarity of transcriptional profiles (initially measured by pairwise correlation) to constrain a low-rank matrix factorization model aimed at predicting deviations from expected fitness when perturbations are combined (fig. S11A; Methods). We then predicted unobserved GIs using this model trained on different fractions of randomly sub-sampled interactions (Fig. 7A; Methods). At 10% sampling, the end result (Fig. 7, B and C) preserves much of the large-scale structure of the map, and the rank order of all interactions is reasonably well-preserved (Spearman ρ = 0.5; Fig. 7D). The model is substantially better than random sampling at predicting the top 10% of interactions (Fig. 7E), preserves pairwise distances among GI profiles (cophenetic correlation 0.7–0.8 at 15% sampling; Fig. 7F and fig. S11B). Notably, the use of Perturb-seq-derived side information significantly improved performance (compare to Figs. S11E-G).

**Figure 7.**
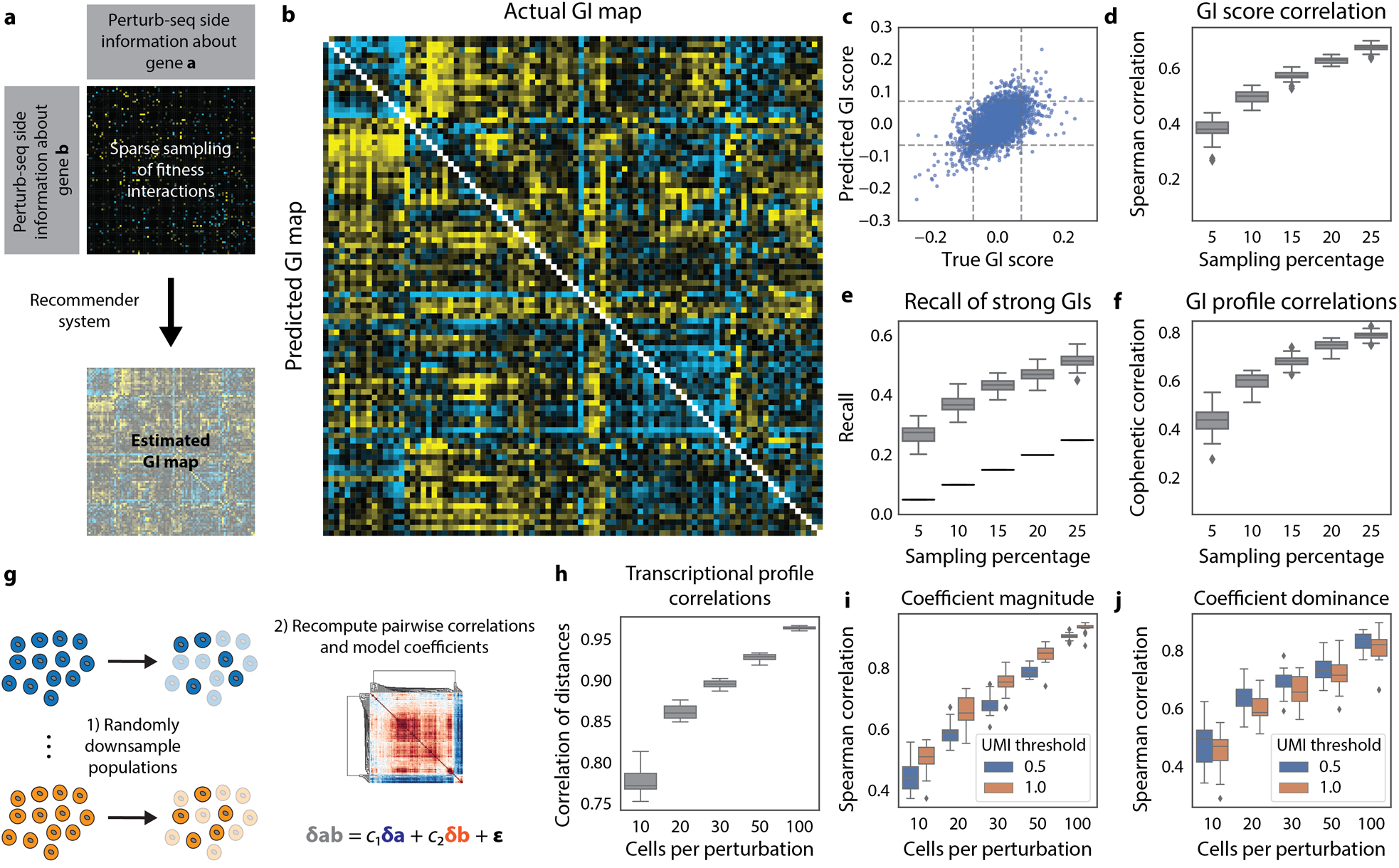
A recommender system enables prediction of fitness GIs. **(A)** Schematic of prediction strategy. Fitness phenotypes of a limited subset of GIs are measured. Each single gene is characterized by its Perturb-seq transcriptional profile, and similarity among these profiles is used as side information to constrain a recommender system model to impute remaining fitness GIs. **(B)** True vs. predicted GI map obtained by prediction from 10% of randomly sampled fitness-level GIs. **(C)** Scatter plot of true and predicted GI scores from (B). The dashed lines show 5% and 95% quantiles, here used to designate strong GIs. **(D)** Spearman correlation between true and predicted GIs at different levels of random sampling of measured GIs. Fifty random subsets were measured for each sampling level. **(E)** At each sampling level, recall denotes the fraction of strong GIs that are correctly scored as strong in the predicted GI map as in panel (C). **(F)** Cophenetic correlation as a function of sampling level, measuring the similarity of correlation structure in the true and predicted GI maps. **(G)** To assess robustness, random subsets of cells bearing given genetic perturbations were randomly sampled at different levels of representation. Analytical results were recomputed across many iterations and compared to the results obtained using all cells. **(H)** Cophenetic correlation between downsampled and true transcriptional profiles used to construct the GI manifold visualization of Figure 3. **(I)** Downsampling analysis of the magnitude of the model coefficients used in Figure 4 and 5, which can be improved by fitting to genes expressed at higher average UMI counts. **(J)** Equivalent analysis for the dominance of model coefficients, used to orient buffering interactions.

Ultimately, predicting novel GIs is difficult precisely because strong GIs deviate from the average interaction behavior. In contrast to model performance for GIs, for example, we observed high predictive power for raw fitness (Spearman ρ ∼ 0.96 at 10% sampling, fig. S11C). By using biased sampling and different side information we could modestly improve GI prediction performance (fig. S11D; Methods). Moving forward, strategies for each of these steps could be computationally learned in larger data sets, further enhancing predictive power.

Finally, scaling issues can also be addressed by measuring fewer cells per perturbation, so that more GIs can be measured in a single experiment. To model this approach, we down-sampled our measured perturbations (gathered at a median of 273 cells per condition) and re-performed some of our general analyses. The correlation distances used to organize perturbations in Figure 3 and the coefficients in our linear model of interaction were both quite stable to downsampling (Fig. 7, G to J), suggesting as few as 50 cells per perturbation could be sufficient. Moreover, stability can be improved by performing model fits using only more highly-expressed genes (Fig. 7, I and J; Methods).

## Discussion

Understanding how genes interact is key to understanding how complex biological phenomena arise from a finite number of genes. To address this challenge, we leveraged both a high-density genetic interaction map, which allows many candidate interactions to be systematically and rapidly profiled (*18*), and rich phenotyping via Perturb-seq, which provides high-dimensional measures of their phenotypic consequences at single-cell resolution (*29*). This synthesis had two direct advantages over approaches that rely solely on univariate measures such as growth. First, transcriptional phenotypes provide specific insights into the underlying biological mechanism of action. Second, it provides a more natural setting to consider how genes interact to generate complex phenotypes: a cell’s state can be thought of as a point on a high-dimensional manifold defined by its transcriptional profile. Genetic perturbations push cells in different directions along this manifold, and interactions emerge when, for example, combinations of perturbations push cells more (SSL) or less (buffering) than expected when combined.

Three broad principles emerge from this approach. First, in fitness-based GI screens information is inevitably lost when genetic interactions are categorized by the umbrella terms “buffering” or “synthetic lethal”—in essence, this implements a type of dimensionality reduction that irreversibly compresses all possible phenotypes onto a line, thereby conflating very different functional relationships. Our geometrically-motivated model of interaction achieves a similar goal but retains more information: instead of being defined by a single number, each interaction is defined by a quantitative model of how the two single perturbations combine. Despite the diversity of our perturbations, this simple linear model fits well (mean R^2^ score=0.71), and yields useful, actionable insights. For example, we could distinguish classes of behavior such as epistasis, suppression, and neomorphism, and order genes into linear pathways. Potential practical applications of this advance include the comprehensive search for true molecular suppressors of disease-causing gene alleles.

Second, unbiased measurements of combinatorially perturbed cells yield direct insight into the mechanisms underlying genetic interactions. For instance, we observed differentiation to multiple distinct cell types, cell cycle arrest, and apoptotic cell death as explanations for synthetic growth defects. Notably, our unbiased activation screen reidentified many of the canonical transcriptional regulators of hematopoiesis, and importantly provided a quantitative description of the regulatory relationships among them. We also identified completely unexpected, widely expressed factors such as calponin (*CNN1*) with equally strong primary and synergistic effects on erythroid differentiation. As rich phenotyping via Perturb-seq enables screen designs that are sensitive to multiple outcomes or with incomplete penetrance, it is a natural strategy to pursue combinatorial searches for factors driving (trans)differentiation, or for genes modifying complex phenotypes that cannot be encapsulated by single markers (*5*, *27*).

Finally, single-cell rich phenotyping methods have the potential to transform the study of complex genetic traits. Recent advances (*33*) now make these methods both highly scalable and highly informative. Combinatorial interaction spaces quickly grow to an intractable size. With Perturb-seq, each cell serves as an individual experiment, allowing facile multiplexing of hundreds of thousands of rich phenotypes at a time. These high-dimensional measurements provide a handle to apply more powerful machine learning methods (*62*). We showed that the GI map underlying a complex trait, fitness, can be computationally predicted from far fewer measurements. This example highlights the possibility of using similar approaches to understand disease-relevant traits. Further, it identifies the challenges and opportunities moving forward: can better ways to compare genetic perturbations (i.e., a data-driven metric on the GI manifold) be learned and can single-cell phenotypic heterogeneity, as seen in the *DUSP9*-*MAPK1* interaction, be exploited to improve predictive power?

Critically, our framework for probing and understanding genetic interactions is compatible both with the many emerging tools for perturbing the state of individual cells— genome editing, base editors, epigenome editing—and the many tools for measuring it— transcriptome, chromatin state, proteome, imaging etc. (*63*). The power of these tools is further enhanced by new computational methods that allow improved design of experiments, such as recommender systems (*62*) and compressed sensing(*64*, *65*). Finally, approaches leveraging allelic series of sgRNAs with different inhibition/activation levels (*35*), or with paired CRISPRi/a systems to look for suppressive interactions (*19*) may be able to exploit single-cell phenotypic heterogeneity to more finely dissect gene-level perturbation responses at scale. The promise of such an approach is exemplified by our analysis of *DUSP9*/*MAPK1* as robust suppressors. Ultimately, the many diverse cell types and cellular behaviors we observe are the product of a finite number of genes working together. By intelligently measuring and exploring the genetic interaction manifold, we can start to create a global view of the nonlinear mapping between genotype and phenotype.

## Acknowledgements

We thank Britt Adamson, Owen Chen, Min Cho, Jacob Corn, Craig Forester and Jeffrey Hussmann, Sherman Weissman and members of the Weissman and Gilbert labs for discussion or unpublished reagents.

## Funding

This work was funded by grants from the National Institutes of Health (NIH, P50 GM102706, U01 CA168370, R01 DA036858 to JSW and NIH, NIH/NCI K99/R00 CA204602 and DP2 CA239597 to LAG). JSW is a Howard Hughes Medical Institute Investigator. T.M.N. is a fellow of the Damon Runyon Cancer Research Foundation DRG-[2211-15]. M.A.H. is a Byers Family Discovery Fellow and is supported by the UCSF Medical Scientist Training Program and the School of Medicine. Oligonucleotide pools were courtesy of the Innovative Genomics Institute.

## Author Contributions

TMN, MAH, LAG, and JSW were responsible for the conception, design, and interpretation of the experiments and wrote the manuscript. MJ, JMR provided critical feedback on the manuscript. LAG, MJ, AX and JMR constructed GI vectors, cDNA vectors and libraries. TMN, JMR, AX, AYG, MAH and LAG conducted the GI mapping and Perturb-seq experiments. MAH performed the GI data analysis. TMN performed the Perturb-seq analysis, GI prediction analysis and modeling of transcriptional genetic interactions. LAG, TMN, AYG and AX designed and conducted validation experiments.

## Competing Interests

TMN, MAH, LAG and JSW have filed patent applications related to CRISPRi/a screening, Perturb-seq and GI mapping. JSW consults for and holds equity in KSQ Therapeutics and Maze Therapeutics. TMN and MAH consult for Maze Therapeutics.

## Data and materials availability

All data is available in the manuscript or the supplementary materials.

---

**Figure S1.**
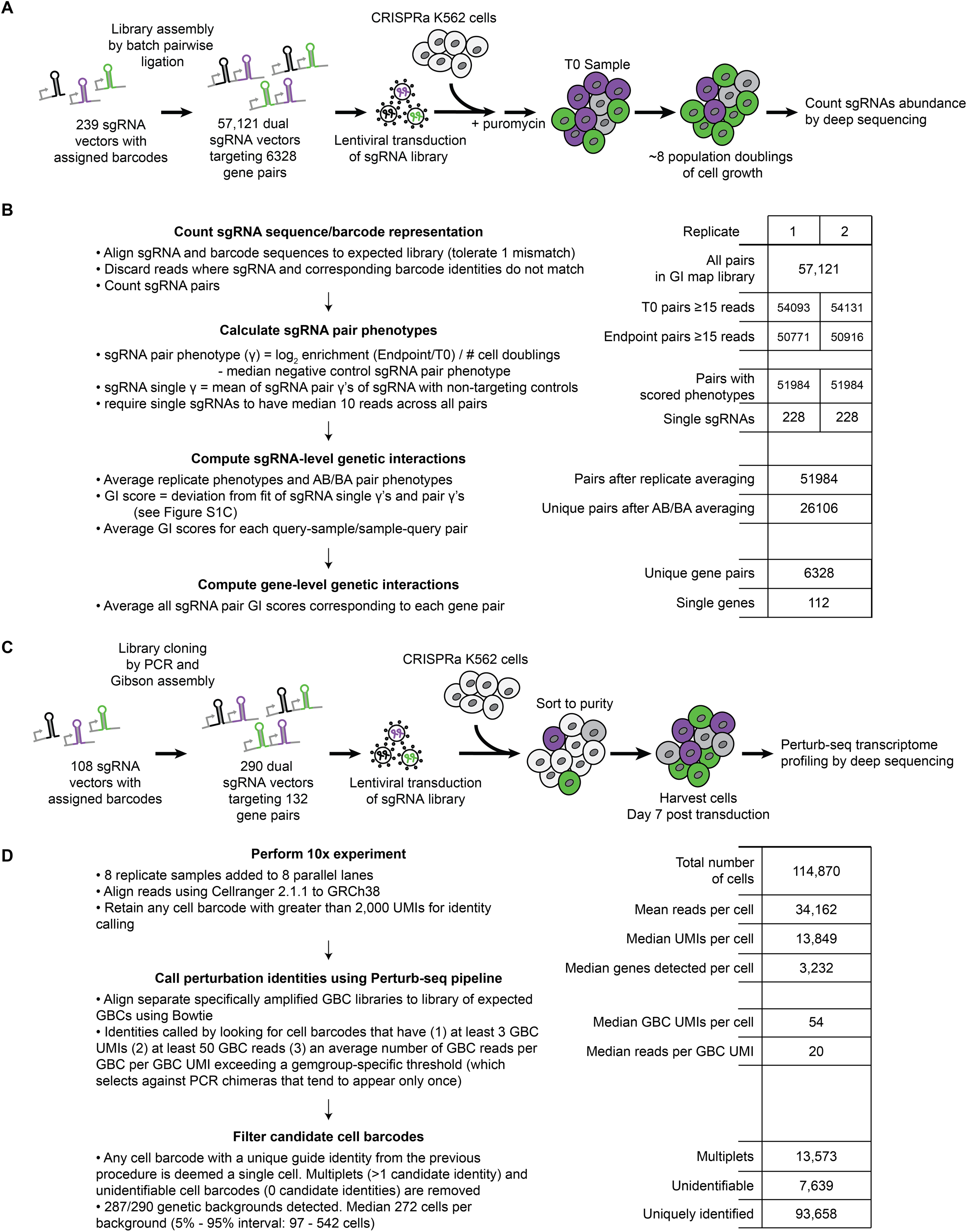
Technical details of GI map and Perturb-seq experiments. **(A)** Schematic of GI map library construction and screen protocol. **(B)** GI map analysis pipeline and statistics at key steps of the pipeline. **(C)** Schematic of Perturb-seq library construction and screen protocol. **(D)** Perturb-seq analysis pipeline and statistics at key steps of the pipeline.

**Figure S2.**
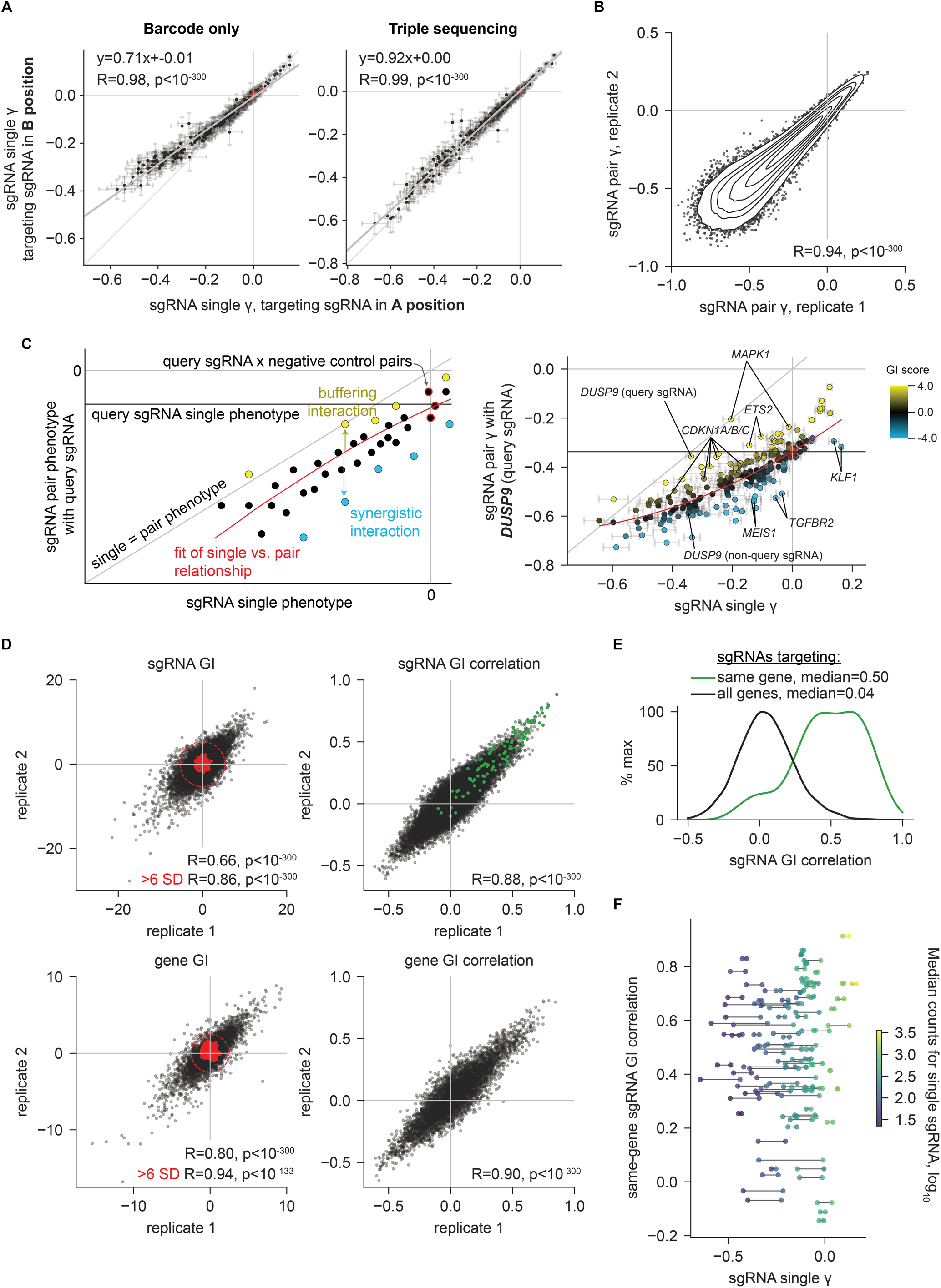
Analysis of sgRNA- and gene-level phenotypes and GIs. **(A)** sgRNA single phenotypes (γ) for sgRNAs in the A or B position in the dual sgRNA expression vector. Points represent the mean and standard deviation of a given sgRNA paired with all non-targeting sgRNAs in the opposite position. sgRNA pair read counts were measured by aligning barcodes only (left) or by aligning both sgRNA positions and barcodes (“triple sequencing,” right). **(B)** sgRNA pair phenotypes (γ) from independent replicates. Contours correspond to 99th, 95th, 90th, 75th, 50th, and 25th percentiles of data density. **(C)** Strategy for calculating sgRNA GIs. (Left) Schematic depicting single sgRNA phenotypes versus sgRNAs paired with a query sgRNA. GIs are calculated from the deviation from the average trend as determined by quadratic fit. (Right) Single versus double phenotypes and sgRNA GIs for a query sgRNA targeting *DUSP9* (DUSP9_+_152912564.23-P1). **(D)** sgRNA- and gene-level GIs and GI profile correlations from replicates. Red points indicate non-targeting control sgRNA pairs and dashed line indicates a radius of 6 standard deviations from non-targeting controls. sgRNA GI correlations are as in Figure 1B. **(E)** sgRNA GI correlations from replicate-averaged screens. Smoothed histograms of all pairs of sgRNAs or only sgRNA pairs targeting the same gene were generated with Gaussian kernel density estimation. **(F)** Relationship between sgRNA single phenotypes and same-gene sgRNA pair GI correlation. Points represent sgRNAs with lines connecting sgRNAs targeting the same gene. Points are colored according to the median read count at screen endpoint across pairs with that sgRNA in the A position.

**Figure S3.**
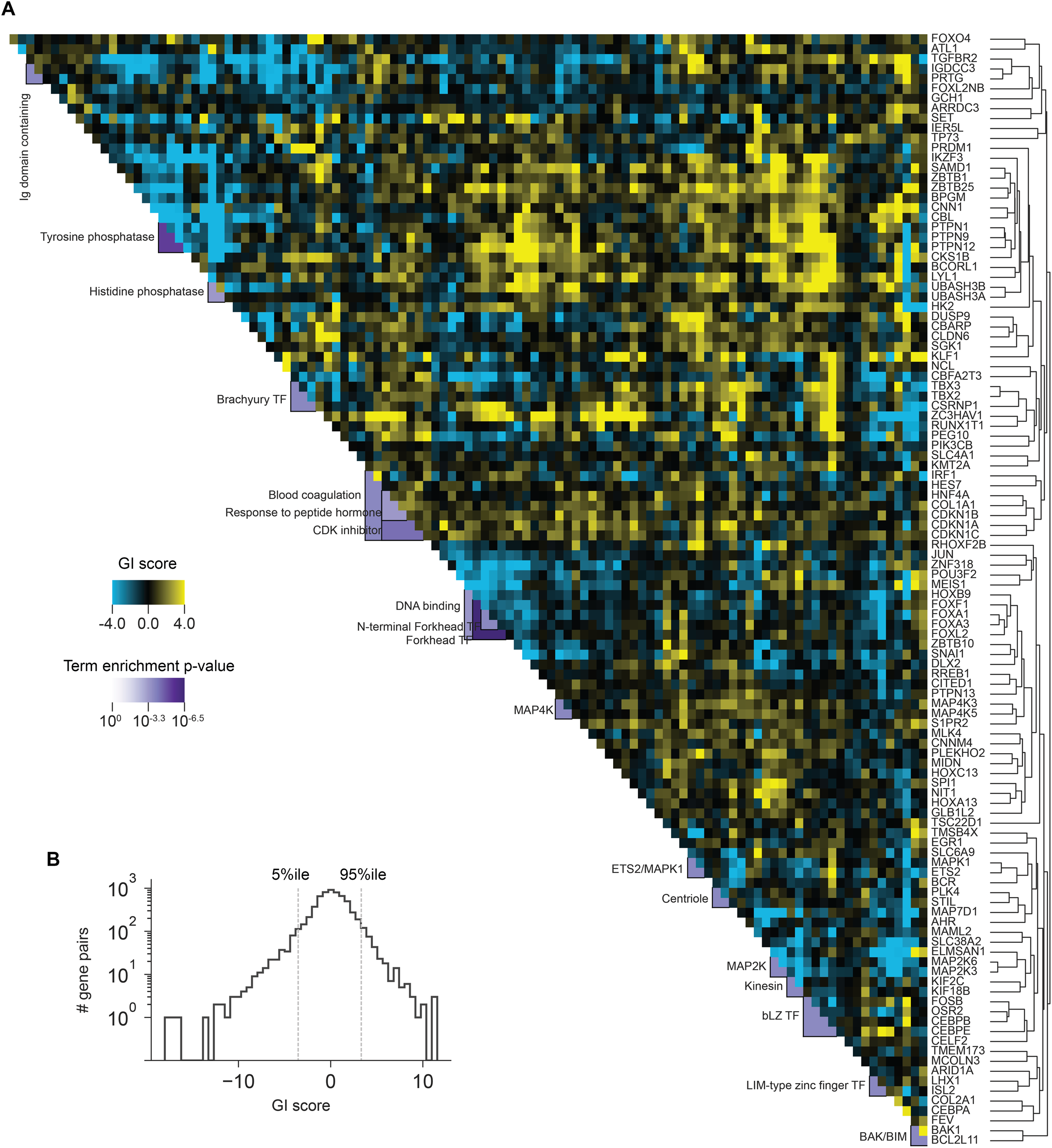
Labeled CRISPRa fitness-level GI map. **(A)** As in Figure 1C, with gene labels along y-axis. **(B)** Histogram of all gene-level GI scores.

**Figure S4.**
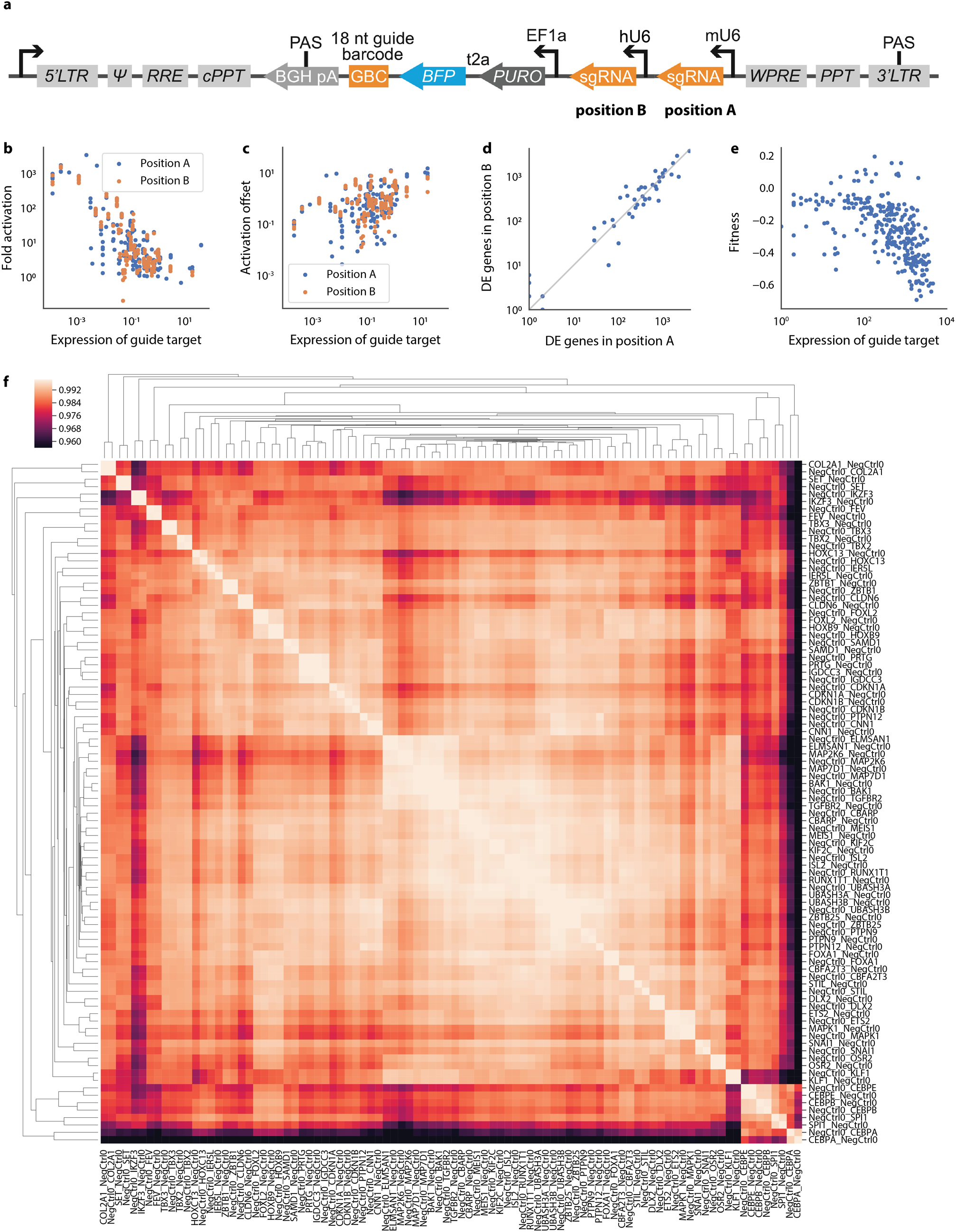
Perturb-seq vector performance. **(A)** Schematic of Perturb-seq dual sgRNA expression vector used for targeting pairs of interacting genes. **(B)** Comparison of target activation to target expression in control cells containing non-targeting sgRNAs. Each dot corresponds to an sgRNA in either the A (blue) or B (orange) position of the vector. **(C)** Equivalent analysis as in (B) but comparing the mean expression difference (in UMI) between cells containing targeting sgRNAs to control cells. **(D)** Comparison between the number of differentially expressed genes (as reported by a Kolmogorov-Smirnov test) when a given sgRNA is in the A or B position of the vector. **(E)** Relationship between fitness (from GI map) and number of differentially expressed genes (as reported by a Kolmogorov-Smirnov test) observed in Perturb-seq experiment for each perturbation. **(F)** Pairwise correlation of expression profiles for all perturbations containing a single targeting sgRNA (in either A or B position) and one non-targeting sgRNA. Presented to assess similarity between phenotypes induced by sgRNAs when present in different positions.

**Figure S5.**
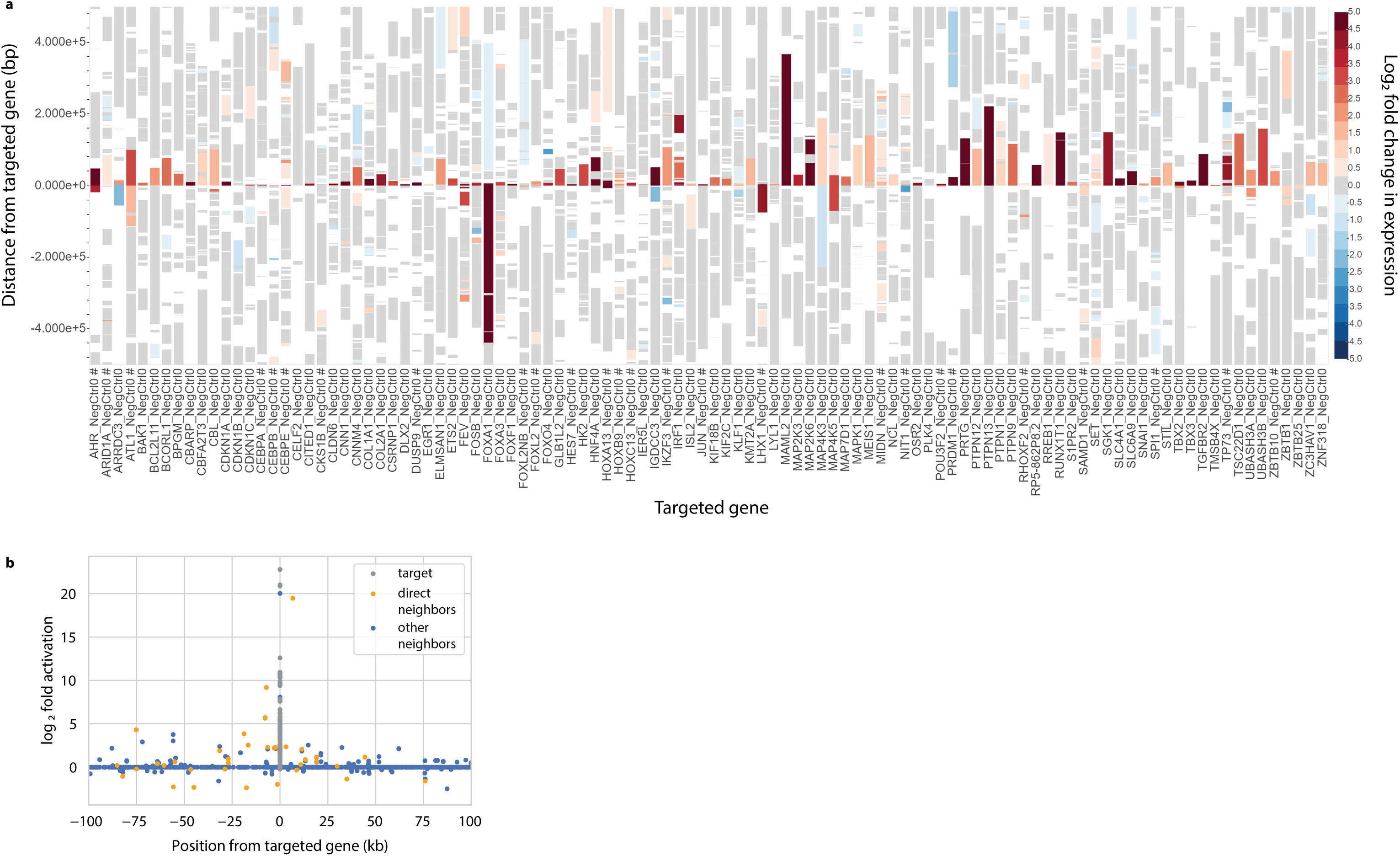
Assessing off-target effects near targeted genes. **(A)** The plot shows expression of transcripts in the neighborhood of the target of each sgRNA. Color indicates the fold change in expression, with gray indicating that expression was indistinguishable from that in control cells containing non-targeting sgRNAs (bootstrap test, *p* < 0.05). A “#” sign in the *x*-axis titles indicates perturbations that induce large changes in total transcriptome size, which may lead to many differentially expressed transcripts that are not the result of off-target effects. **(B)** Plot comparing fold expression change of targeted transcripts (gray), the two transcripts up and downstream of the targeted one (orange), and 19 other neighboring transcripts in each direction (blue) as a function of genomic distance.

**Figure S6.**
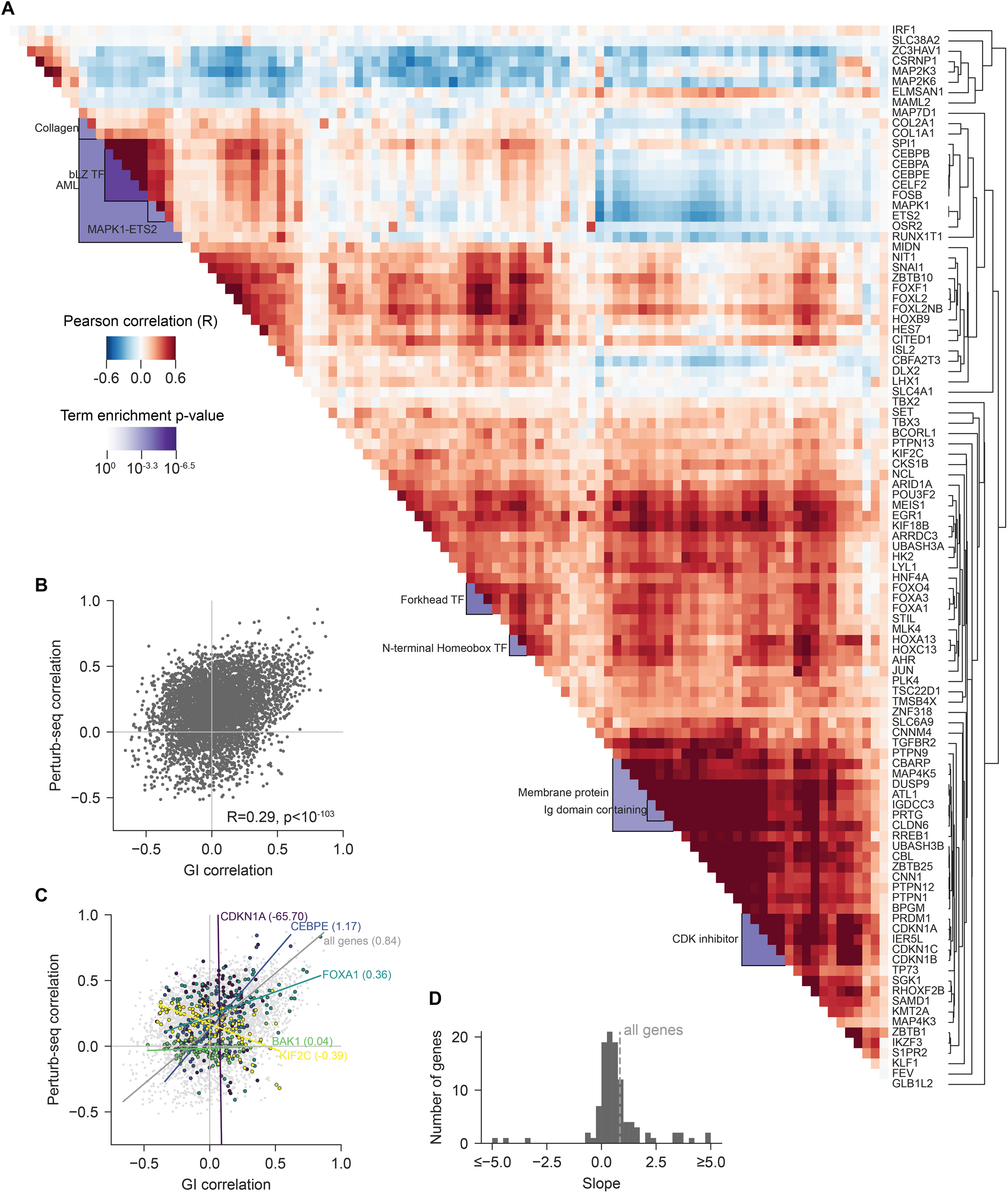
Functional clustering via single sgRNA perturb-seq expression correlation. **(A)** Heatmap of differential expression correlation between pairs of genes upon overexpression with CRISPRa. Pearson correlation was calculated based on variably expressed genes across all perturbations, excluding targeted genes. Hierarchical clustering and cluster annotation were performed as in Figure 1C. **(B)** Relationship between fitness-level GI correlations and perturb-seq differential expression correlations. **(C)** As in B, with gene pairs that include selected genes highlighted. For all genes and for the highlighted genes, line represents the first principal component with the slope in parenthesis. **(D)** Histogram of GI correlation vs perturb-seq correlation slopes for each gene in the map.

**Figure S7.**
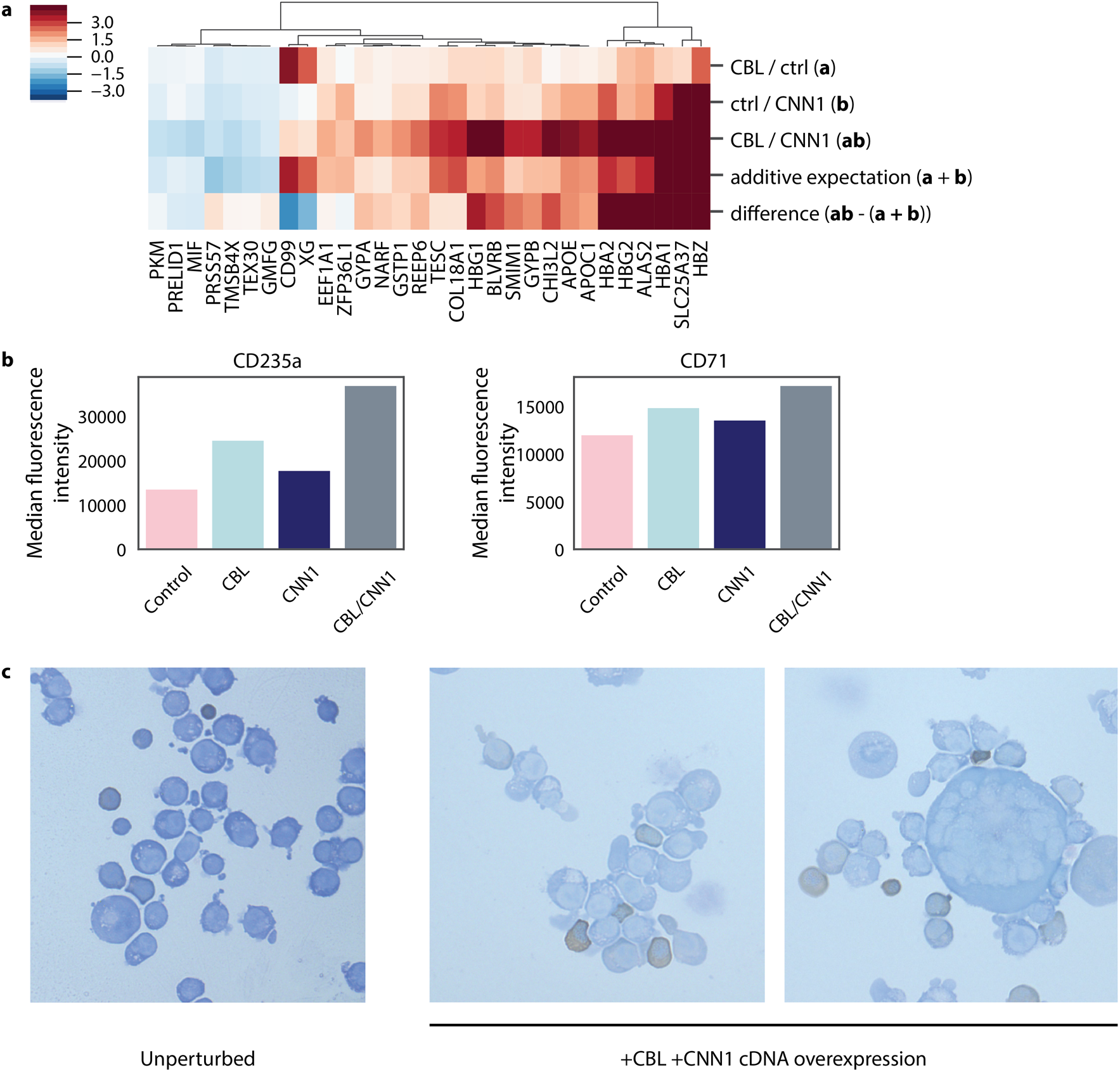
Validating *CBL*/*CNN1* interaction. **(A)** Comparison of expression data from Fig. 2G to an additive model of genetic interaction. **(B)** Median expression of CD235a measured by flow cytometry of antibody-stained K562 cells upon cDNA overexpression of the indicated genes. **(C)** Equivalent analysis for CD71. **(D)** Cytospin images showing benzidine staining of hemoglobin in HUDEP2 cells upon cDNA overexpression of the indicated genes.

**Figure S8.**
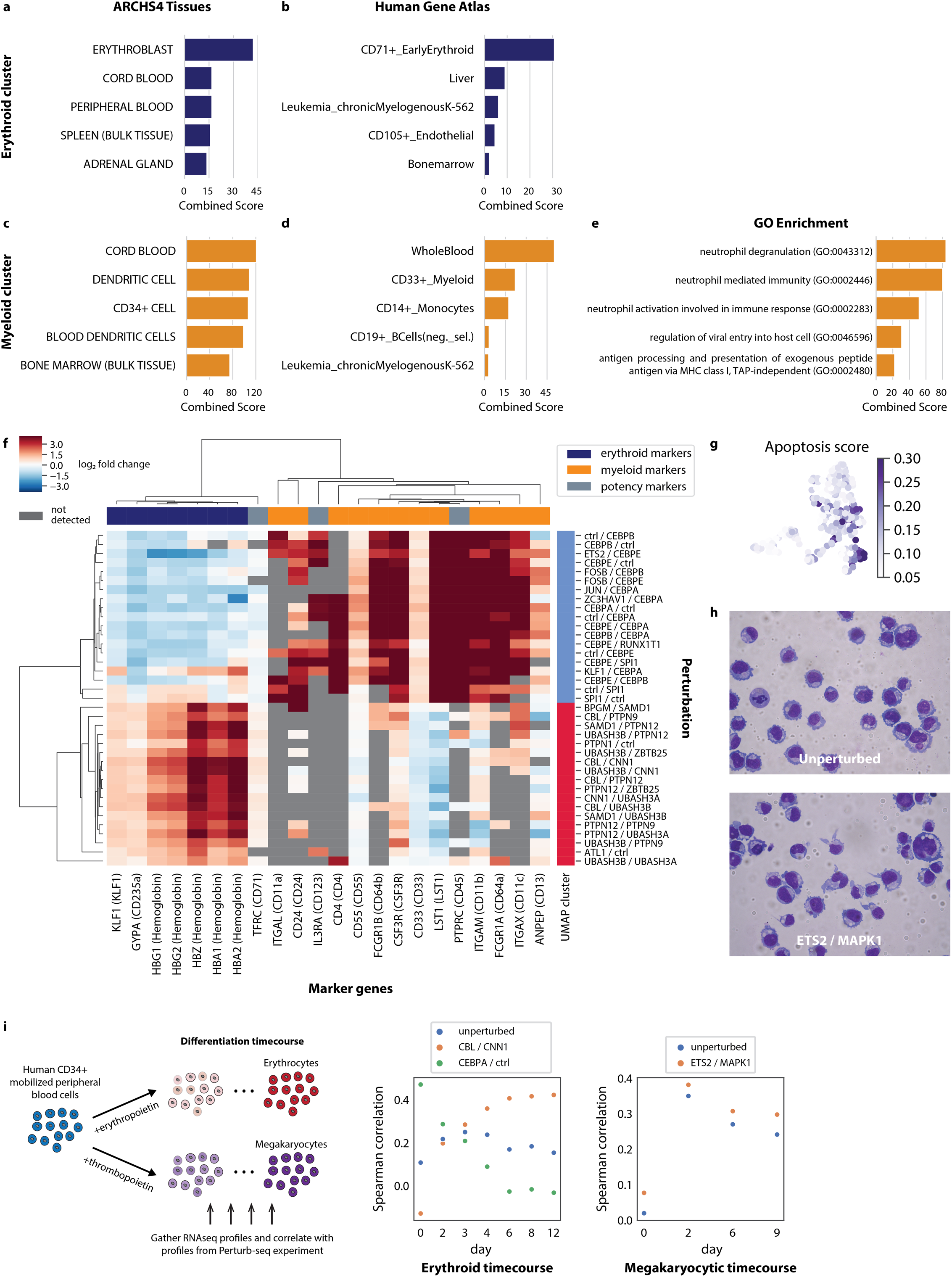
Expression of developmental markers in Perturb-seq experiment. (A)-(E) The panels show term enrichment of genes overexpressed in either the erythroid or granulocyte clusters of Figure 3 within the indicated databases. Analyses were performed using Enrichr (*67*). **(F)** Expression of curated developmental marker genes in perturbations belonging to the erythroid and granulocyte clusters. Color indicates fold change in expression, with gray indicating that a marker was not detected. **(G)** Apoptotic activity within Figure 3. Color is scaled by mean Z-normalized expression of a panel of pro- and anti-apoptotic genes. **(H)** May-Grünwald Giemsa staining of K562 CRISPRa cells expressing either non-targeting sgRNAs (unperturbed) or *ETS2*/*MAPK1*-targeting sgRNAs. **(I)** Differentiation time course experiment. RNAseq profiles of human CD34+ cells induced to differentiate were obtained at the indicated time points. The plots compare how correlated these transcriptional profiles are with the indicated transcriptional profiles of K562 cells in the Perturb-seq experiment.

**Figure S9.**
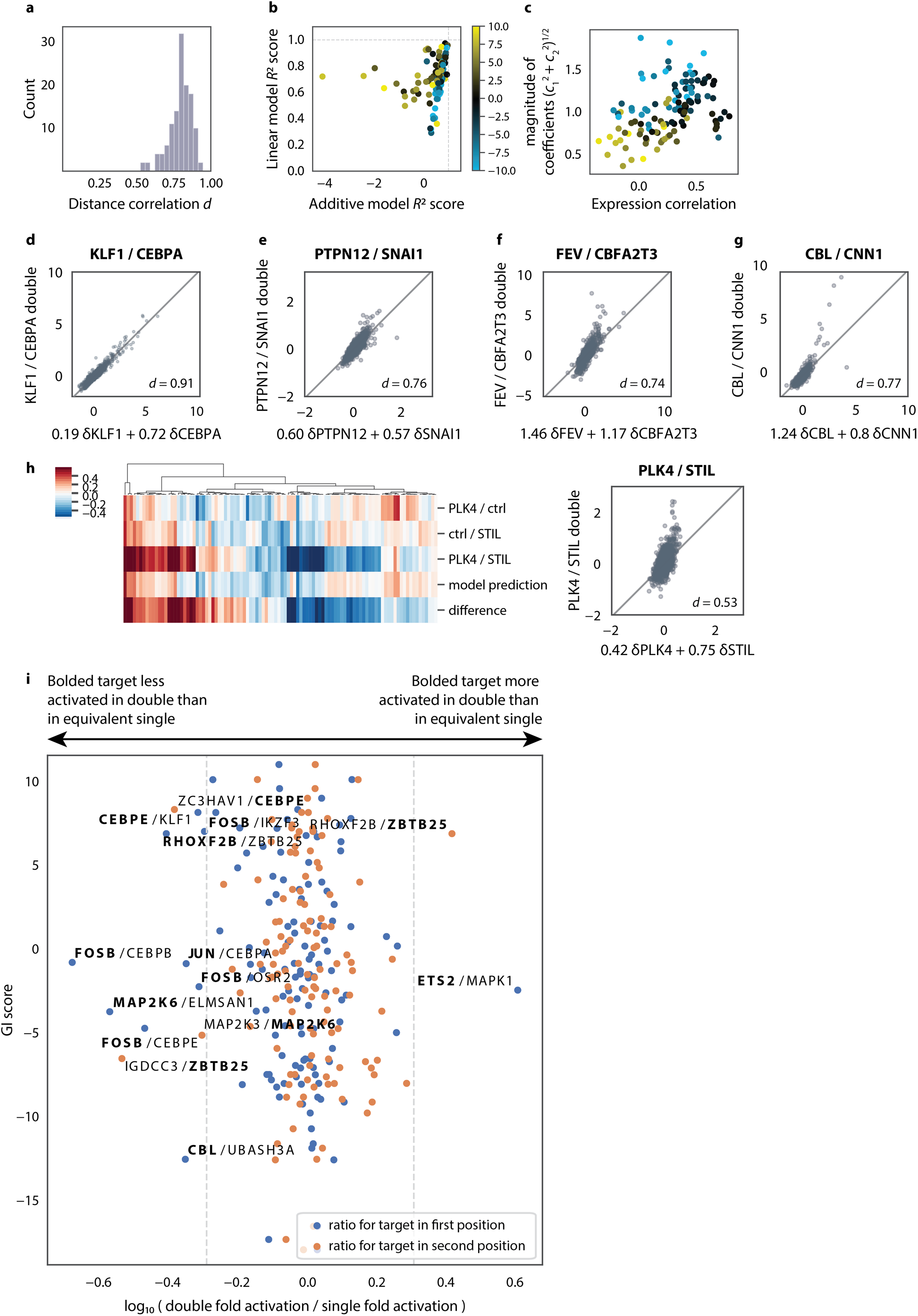
Properties of model of transcriptional genetic interactions. **(A)** Distance correlation between model prediction and true double profile for all perturbations in the Perturb-seq experiment. *d* = *cor* (𝑐_1_**𝛅a** + *c*_2_**𝛅b**, **𝛅ab**). Cf. Figure 4B. **(B)** Comparison of model fit (measured by *R*^2^ score) for the linear model of genetic interactions and a fixed additive model in which **𝛅ab** = 𝛅**a** + 𝛅**b**. **(C)** Comparison between correlation of transcriptional profiles and magnitude of coefficients in the model of genetic interactions. **(D)-(G)** Scatter plots comparing expression in the indicated double perturbation backgrounds and model predictions from the corresponding single perturbations. Each dot denotes a gene used in fitting the model. Models are fit by robust regression, so outlier genes are censored during fitting. **(H)** Perturb-seq dissection of the neomorphic *PLK4*/*STIL* combination. Cf. Figure 2. **(I)** Analysis assessing the extent to which genetic interactions may derive from direct activation of one interaction partner by the other. If the target gene (bolded) is more expressed in a given double perturbation (e.g. *ETS2*/*MAPK1*) than in the equivalent single (e.g. *ETS2/*control), it will lie to the right, and vice versa. Blue dots make the comparison for targets in the first (A) position of the vector, and orange dots in the second (B) position.

**Figure S10.**
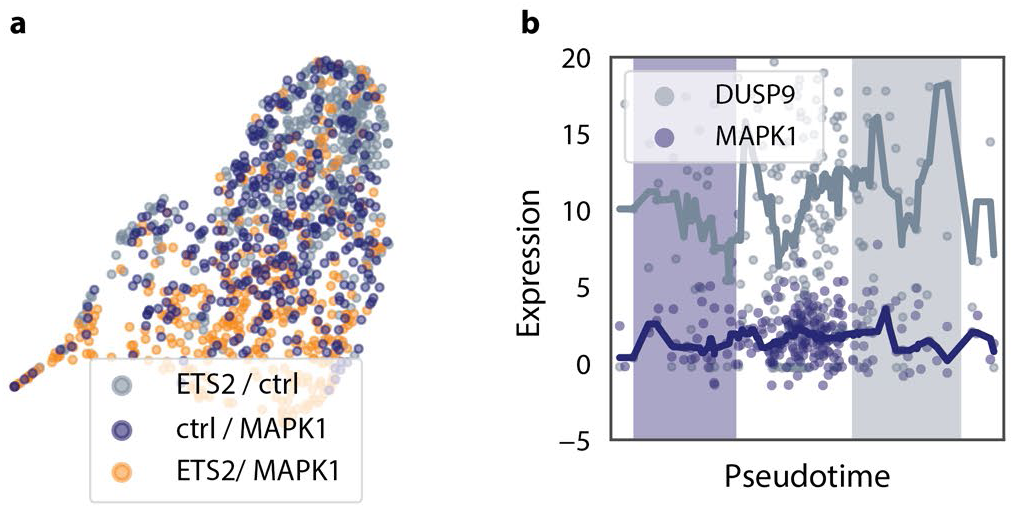
Single-cell trajectories of GIs. **(A)** UMAP projection of single cells (dots) involved in the *ETS2*/*MAPK1* interaction, which largely recapitulate the average behavior. Cf. Figure 4F. **(B)** Expression of *DUSP9* and *MAPK1* in *DUSP9*/*MAPK1*-perturbed cells (dots) ordered by pseudotime. The lines show median-filtered expression.

**Figure S11.**
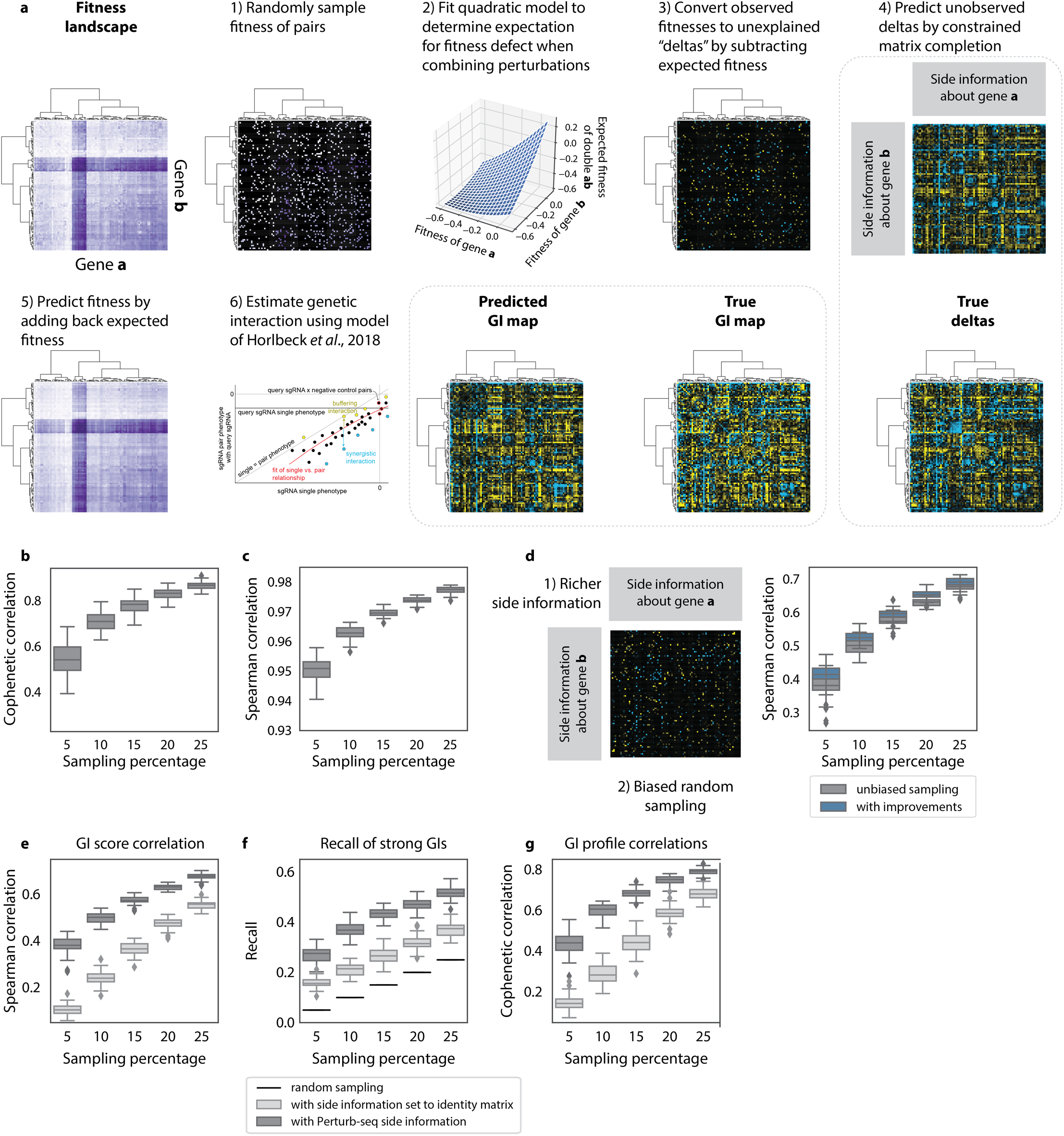
Approach to predicting fitness-level genetic interactions. **(A)** Schematic of approach. Each step is explained in detail in Methods. **(B)** Cophenetic correlation as a function of sampling level, measuring the similarity of correlation structure in the true and predicted GI maps. This figure uses Euclidean distance instead of cosine distance, cf. Figure 7F. **(C)** Spearman correlation between true and predicted fitness phenotypes at different levels of random sampling. Fifty random subsets were measured for each sampling level. **(D)** Improving prediction performance through altered sampling and side information. A biased strategy to sample interaction pairs and modified side information (see Methods) were used as inputs to prediction. The figure compares performance to the results from Figure 7D (“unbiased sampling”). **(E)-(G)** The recommender system can also be used in the absence of side information to perform inference. The figures compare performance with and without Perturb-seq side information. Compare with Figs. 7D-F.

## Supplementary Tables

**Table S1.** Genes targeted in CRISPRa GI map.

**Table S2.** sgRNAs and barcodes in CRISPRa GI map library.

**Table S3.** GI map sgRNA pair read counts and phenotypes.

**Table S4.** GI map sgRNA-level GIs and GI correlations.

**Table S5.** GI map gene-level GIs and GI correlations.

**Table S6.** DAVID annotations for clusters annotated in (A) GI map and (B) Perturb-seq correlation map.

**Table S7.** Perturb-seq experiment statistics for each perturbation.

**Table S8.** Effects of CRISPRa on neighboring genes.

**Table S9.** Fit statistics for the GI manifold linear model by perturbation.

## Experimental Methods

### Cell Lines and Lentivirus

K562, HEK293 and HUDEP2 cell lines were cultured at 37°C 5% CO2 in standard cell culture incubators. K562 (female) cells were grown in RPMI-1640 with 25mM HEPES and 2.0 g/L NaHCo_3_ in 10 % fetal bovine serum (VWR Seradigm), 2 mM glutamine (ThermoFisher), 100 units/mL streptomycin and 100 µg/mL penicillin (ThermoFisher). HEK293T (female) cells used for packaging lentivirus were grown in Dulbecco’s modified eagle medium (DMEM) in 10 % fetal bovine serum (VWR Seradigm), 100 units/mL streptomycin and 100 µg/mL penicillin with 2mM glutamine (ThermoFisher). HUDEP2 cells were cultured in SFEM (StemCell Technologies) with 1µM dexamethasone, 1µg/ml doxycycline, 50ng/mL hSCF (Peprotech), 50ng/mL EPO (Peprotech) and 100 units/mL streptomycin and 100 µg/mL penicillin (*52*). Lentivirus was produced by transfecting HEK293T with standard packaging vectors using *Trans*IT®-LTI Transfection Reagent (Mirus, MIR 2306). Viral supernatant was harvested 72 hours following transfection and filtered through a 0.45 μm PVDF syringe filter. We used our previously published CRISPRa K562 cell line for all CRISPRa experiments. Briefly, to construct the CRISPRa K562 cell line, we lentivirally transduced K562 cells to stably express scFV-sfGFP-VP64, rtTA and a doxycycline inducible dCas9-10xGCN4-P2A-mCherry construct (*39*). We added doxycycline to activate dCas9-10xGCN4-P2A-mCherry expression then sorted the CRISPRa K562 cells by flow cytometry using a BD FACS Aria2 for stable co expression of mCherry and GFP signal which marks expression of dCas9-10xGCN4-P2A-mCherry and the scFV-sfGFP-VP64 proteins respectively. We plated this polyclonal CRISPRa population to single cell clones and characterized CRISPRa activity as described (*39*) to generate our CRISPRa K562 cell line. K562 are originally obtained from ATCC. HUDEP2 are a gift from Jacob Corn. All cell lines were routinely tested for mycoplasma (MycoAlert, Lonza).

### CRISPRa, sgRNA and cDNA plasmids

We used previously described lentiviral vectors to express the CRISPRa scFV-sfGFP-VP64, rtTA and a doxycycline inducible dCas9-10xGCN4-P2A-mCherry constructs (*39*). The sgRNA constructs and libraries used for GI mapping and perturb-seq experiments are described in detail in the GI and perturbs-seq sections of the methods below. To confirm individual GI and manifold phenotypes by CRISPRa we used two sgRNA vectors that each encodes a modified mouse U6 promoter that drives expression of an optimized *S. pyogenes* sgRNA constant region (*68*) as well as either GFP or BFP and a puromycin resistance cassette separated by a T2A sequence from an Ef1Alpha promoter. For cDNA experiments that confirm manifold phenotypes, we cloned each specified cDNA into the LeGo iG2 and iC2 cDNA overexpression vectors by blunt ligation (*69*).

### GI dual sgRNA library plasmid design and construction

The GI dual sgRNA library vector is a previously published sgRNA lentiviral plasmid (*18*). In the final GI library sgRNA vector, modified mouse and human U6 promoters express the 5’ and 3’ sgRNAs respectively. Each sgRNAs encodes the same optimized *S. pyogenes* sgRNA constant region (*68*). To enable measurement of lentiviral vector recombination during reverse transcription by Illumina sequencing, we encoded 4 randomized 16 base pair DNA barcodes in the GI library dual sgRNA vector as previously described (*18*). The GI lentiviral sgRNA construct co-expresses BFP and a puromycin resistance cassette separated by a T2A sequence from an Ef1Alpha promoter.

The GI CRISPRa libraries were prepared by library cloning protocols as previously described (*18*). Briefly, we encoded each sgRNA on two oligonucleotides (Integrated DNA Technologies) which were annealed and ligated into a modified pSICO vector (pLG_GI1) encoding the optimized *S. pyogenes* sgRNA constant and two 16 base pair random DNA barcodes. We sanger sequenced each vector to assign DNA barcodes to each sgRNA in the library and then pooled the library manually. This starting sgRNA library lacks U6 promoters. As previously described, we cloned either a modified human or modified mouse U6 promoter into our pooled sgRNA library by restriction digest with XhoI/BstXI followed by ligation, creating two libraries where each vector encodes 1 U6-sgRNA cassette and 2 unique barcodes (pLG_GI2 and pLG_GI3). Finally, we restriction digested the mouse U6-sgRNA library with AvrII and KpnI and the human U6-sgRNA library with XbaI and KpnI, we gel isolated the appropriate fragments of DNA and ligated these two libraries together creating our final GI sgRNA library vector. This GI dual sgRNA library encodes 2 sgRNAs with the 5’ sgRNA expressed from the mouse U6 promoter and the 3’ position from the human U6 promoter as well as the 4 unique DNA barcodes (pLG_GI4) as previously described (*18*).

### CRISPRa GI screening

CRISPRa K562 cells were infected with CRISPRa GI dual sgRNA libraries by spinoculation for 2 hours at 1000g in the presence of 8 μg/mL polybrene (*35*). The lentiviral infection was scaled such that on average each cell is infected with one GI dual sgRNA library vector as measured by the lentivirally encoded BFP signal. At each point of the CRISPRa GI screen we maintained a library coverage of at least 3000 cells per GI perturbation except at the initial infection where we infected 250 cells per sgRNA. Two days after lentiviral infection, cells were selected with 0.9-1 μg / mL puromycin (Sigma) for 2 days, and recovered with addition of fresh media for a ∼48 hour recovery. For the screen, populations of K562 cells expressing the CRISPRa GI library were harvested at the outset of the experiment or after ∼8-9 population doublings. Two biological replicates of the GI screen were performed. Genomic DNA was harvested from each sample; the sgRNA and barcode encoding regions were amplified by PCR and then sequenced at high coverage as previously described (*18*).

### Perturb-seq plasmid design and construction

To generate our library of dual sgRNA expression Perturb seq vectors, we began by cloning sgRNAs into single sgRNA expression vectors. sgRNAs destined for position A of the dual sgRNA vector were cloned into pBA439 (Addgene, #85967), which expresses sgRNAs with constant region 1 from an mU6 promoter, while sgRNAs destined for position B were cloned into pMJ117 (Addgene, #85997), which expresses sgRNAs with constant region 3 from an hU6 promoter. sgRNAs (ordered as annealed oligos from IDT with BstXI/BlpI overhangs) were ligated into parental vectors after vector digestion with BstXI and BlpI. We then performed clonal isolation and Sanger sequencing to establish our library of single sgRNA expression vectors. Next, we prepared inserts and vectors for Gibson assembly. We PCR amplified the sgRNA expression cassettes consisting of a U6 promoter, protospacer, and constant region. pBA439-derived cassettes were amplified with oMJ0571 (GCTGAGTGTAGATTCGAGCAAAAAAAGCACCGACTCG) and oMJ0572 (gaagttattaggtccctcgac) while pMJ117-derived cassettes were amplified with oBA276 (cggtaatacggttatccacg) and oBA281 (GCTCGAATCTACACTCAGC). We Gibson assembled XhoI/HpaI-digested pBA571 (Addgene, pooled library #85968), pBA439-derived cassettes, and pMJ117-derived cassettes to generate dual sgRNA expression vectors. We performed clonal isolation and Sanger sequencing of the two protospacers and corresponding GBCs to establish a library of uniquely barcoded dual sgRNA expression Perturb seq vectors (Table S2)

### Perturb-seq viral production and lentiviral titering

Lentivirus for the 295 dual sgRNA expression Perturb seq vectors was produced in array to prevent sgRNA-barcode uncoupling. To control the representation of sgRNAs in cells at the time of scRNA-seq (7 days after transduction), viral pooling was performed after titering to account for packaging variability and sgRNA effects on cell growth over the course of the experiment. To titer, sgRNA expression constructs were transduced into CRISPRa K562 cells (stably expressing the sunCas9 system (*39*)) via centrifugation (2 hours at 1000 × g). Cells were grown without selection for 7 days and analyzed by flow cytometry on a LSR-II flow cytometer (BD Biosciences). BFP expression was used to gate for transduced cells. The final lentiviral pooling was corrected for both this viral titer and desired representation (for example, negative control vectors were included at 6-fold higher representation to ensure adequate sampling).

### Perturb-seq screening

Our pooled dual sgRNA CRISPRa library was spinfected into CRISPRa K562 cells (*39*) (2 hours at 1000 × g) with a target multiplicity of infection (MOI) of 0.04. Post centrifugation, cells were transferred to a spinner flask for growth. After 3 days of growth, we measured (by BFP expression) an MOI of ∼0.03 and transduced cells were sorted to near purity on a BD FACSAria2. Cells were maintained at >88% viability and >94% GFP+BFP+ over the course of the experiment, indicating stable CRISPRa and sgRNA vector expression. On day 7 post-transduction, cells were separated into droplet emulsions using the Chromium Controller across 8 lanes of the Chromium Single Cell 3’ Gel Beads v2 (10x Genomics). Cells were loaded to recover ∼13,000 cells per lane at an estimated coverage of ∼350 cells per sgRNA.

### Library preparation and sequencing

The scRNA-seq library for our GBC Perturb-seq screen was prepared according to the Chromium Single Cell 3’ Reagent Kits v2 User Guide (10x Genomics CG00052). Library molecules containing guide barcodes (GBCs) were specifically amplified using KAPA HiFi ReadyMix with 180 ng of the final library as template and 0.6 mM of the custom P5 primer 052-P5 (5’-AATGATACGGCGACCACCGAGATCTACAC-3’) and 0.6 mM of custom i7-barcoded specific amplification primers (table below). PCR cycling was performed according to the following protocol: (1) 95 C for 3 min, (2) 98 C for 15 s, then 70 C for 10 s (14 cycles) (3) 72 C for 1 min. The resulting GBC sequencing library was purified via a 0.8X SPRI selection followed by selection of fragments length 350-525 bp using the BluePippin (Sage Science).

All scRNA-seq and GBC sequencing libraries were sequenced on 1 lane of a NovaSeq S2. The libraries were sequenced with a 26 bp Read 1, 90 bp Read 2, and 8 bp Index Read 1.

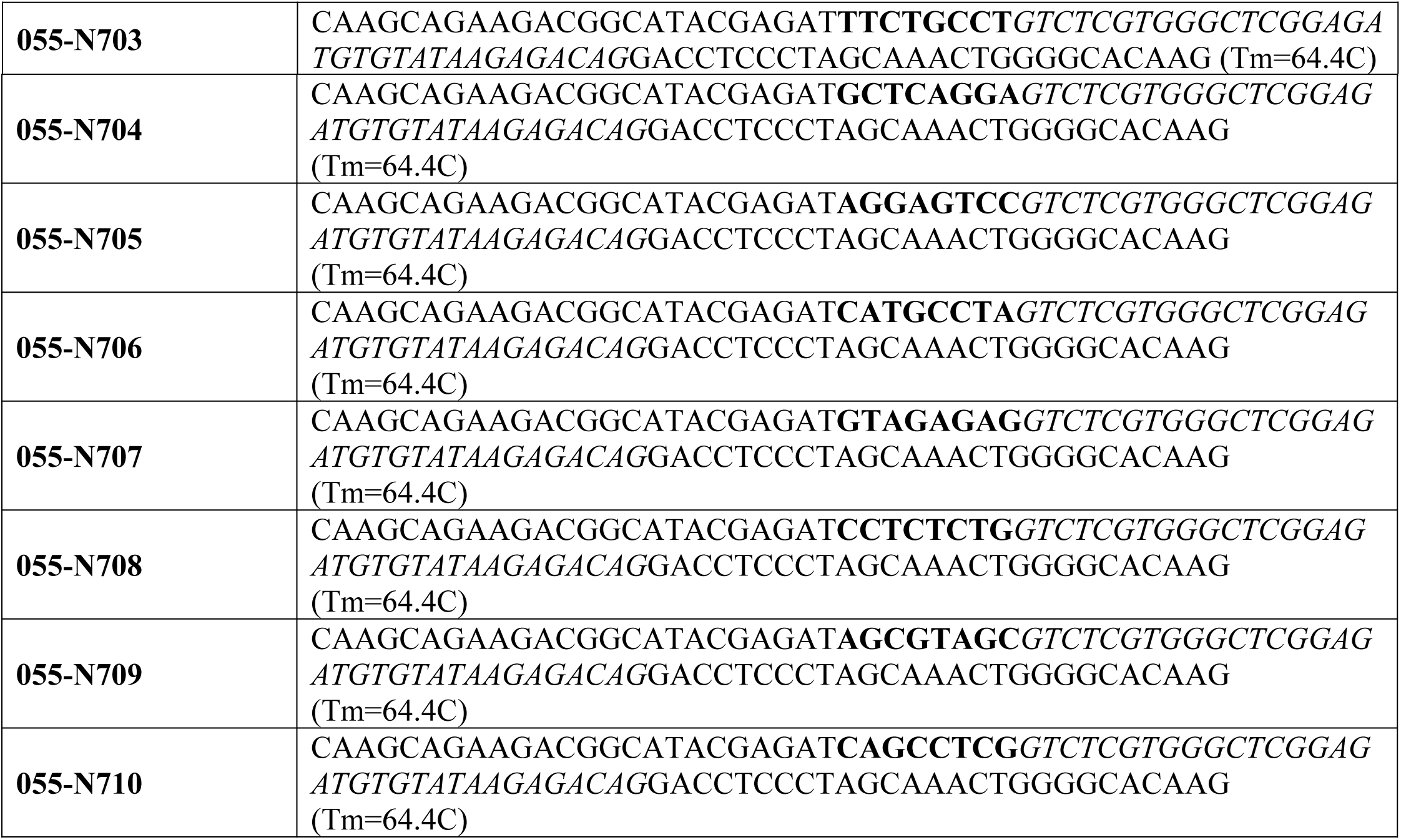

### CRISPRa and cDNA validation experiments

Individual phenotype re-test experiments for sgRNA or cDNA pair phenotypes from the GI and perturb-seq screens were performed as pure population assays or as dual color (BFP/GFP) immunophenotyping experiments on a partially transduced population of CRISPRa K562, K562 or HUDEP2 cells. Briefly, for multiplexed immunophenotyping cells were co-transduced at ∼5-60% infection with two lentiviral vectors marked with either BFP, mCherry or GFP each encoding a single sgRNA or cDNA. This assay enables us to track uninfected cells, cells that express each single sgRNA or cDNA or cells that express a pair of sgRNAs or cDNAs within one internally controlled sample by flow cytometry over time to quantify how each single perturbation or pair of perturbation influences cell phenotypes. Three to six days following infection, cells were counted and stained with APC, APC-Cy7 or PE-Cy7 conjugated antibodies as indicated (BioLegend) and then analyzed by flow cytometry on a BD LSR-II. For hemoglobin and cell morphology assays, infected cell populations were sorted to purity by flow cytometry using a BD FACS Aria2 or a Sony SH800S Cell Sorter for stable BFP, mCherry or GFP signal which marks sgRNA or cDNA expression 2 days following infection and then cultured for additional 3-4 days.

### Cytology

CRISPRa K562 or HUDEP2 cells were concentrated by cytocentrifugation at 400*g* for 8 minutes onto glass slides using a Shandon Cytospin 3 (Thermo Fisher Scientific). Slides were fixed in methanol for 30 seconds, stained in May-Grünwald solution (Sigma-Aldrich) for 10 minutes, and stained in a 1:5 dilution of Giemsa solution (Sigma-Aldrich) in distilled water for 20 minutes. Stained slides were then rinsed in distilled water and allowed to dry before being covered with a coverslip. Cells were imaged on a Zeiss AxioImager M1.

### Hemoglobin expression analysis

Hemoglobin expression was visualized by flow cytometry on a BD LSR-II by staining cells with PE conjugated anti-human HbF antibody as per manufacturers protocol (BD Biosciences). For histochemical analysis of hemoglobin, HUDEP2 cells were stained with *o*-dianisidine. Briefly, cytospun slides were fixed in methanol for 30 seconds, stained in *o*-dianosidine solution (Sigma-Aldrich, 1% w/v in methanol) for 5 minutes, and stained in peroxide solution (1 vol. 30% H_2_O_2_ and 11 vol. 70% EtOH) for 2.5 minutes. Slides were then counterstained with May-Grünwald solution for 10 minutes, rinsed with distilled water, dried, and covered with a coverslip. Cells were imaged on a Zeiss AxioImager M1.

## Computational Methods

### GI map analysis

#### Availability

All analysis of GI map experiments was performed using a custom package built in Python. This package was previously published in part (*18*), but the complete package used in the present manuscript and the associated notebooks (<u>names appear underlined</u>) used to process the data and generate all the figures will be released alongside the paper.

#### GI map gene and sgRNA selection

(GI_library_construction)

Genes with a gene growth phenotype (γ) of absolute value greater than or equal to 0.05 in a CRISPRa v1 growth screen in K562 (Table S1; (*35*)). The gene set was further filtered for genes with a “discriminant score” (the product of the phenotype Z-score and -log(Mann-Whitney P-value)), of 20 or greater in at least one screen in our CRISPRa activity score dataset (*70*). The two sgRNAs targeting each gene with the greatest absolute value growth phenotype were selected, standardizing CRISPRa v1 sgRNA length to G(N19)NGG and excluding sgRNAs containing BstXI, BlpI, SbfI, XhoI, AvrII, and XbaI sites. 16 non-targeting sgRNA controls were randomly selected from CRISPRa v1 with the same length adjustment and restriction site filters. sgRNAs were ordered as pairs of individual oligonucleotides (Integrated DNA Technologies, San Diego, CA) with the following flanking sequences:

Top oligo: TTG + G (N19) + GTTTAAGAGC

Bottom oligo–reverse complement of: CTTGTTG + G(N19) + GTTTAAGAGCTAA sgRNA sequences are provided in Table S2.

#### GI map screen data processing

<u>(GI_data_processing)</u>

##### Sequence alignment

Read alignment was performed as previously described (*18*). Sequencing reads resulting from triple sequencing were obtained as three parallel fastq files representing the upstream and downstream sgRNAs and the reverse-complement of the pair barcode. sgRNAs and barcodes were aligned to the sequences present in the CRISPRa GI map library using custom Python 2.7 scripts (github.com/mhorlbeck/GImap_tools/tripleseq_fastqgz_to_counts.py), allowing for one mismatch in each of the sgRNAs and barcodes. Only reads in which the sgRNAs and barcodes all mapped to the library and matched to the same pair ID were counted for downstream analysis (with the exception of the “barcodes only” analysis in Figure S2A). The barcode did not match the A position in ∼4% of reads and did not match the B position in ∼16% of reads, roughly proportional to their distances in the paired sgRNA vector and consistent with our previous findings (*18*). Raw reads are deposited with GEO (GSENNNNN) and read counts are provided in Table S3.

##### Calculating GI scores

Phenotypes, sgRNA-level GI scores, and gene-level GI scores were calculated essentially as described in (*18*) using custom Python 2.7 scripts (github.com/mhorlbeck/GImap_tools/GImap_analysis.py). sgRNA pair phenotypes were obtained by calculating the log_2_ enrichment of read counts at the endpoint compared to T0, with a pseudocount of 10 and filtering all sgRNAs for which there were fewer than a median of 15 reads across all pairs with that sgRNA in either the A or B position. The log_2_ enrichment values were divided by the number of cell doublings between T0 and endpoint (replicate 1: 8.079, replicate 2: 8.322) to obtain the growth phenotype γ (*71*). Replicate phenotypes were then averaged except for analyses directly comparing replicates. Finally, corresponding AB and BA sgRNA pair phenotypes were averaged. Phenotypes are provided in Table S3.

Single sgRNA phenotypes were calculated from the mean of all sgRNA × negative control pairs. sgRNA-level GI scores were then calculated by fitting a quadratic curve to the relationship of single phenotypes to the sgRNA pair phenotype when paired with a given query sgRNA (Figure S2C). The y-intercept of the quadratic curve was set at the single phenotype for the query sgRNA. For each pair, the GI score equaled the difference between the measured pair phenotype and the fit expectation at the single phenotype, standardized to the standard deviation of the query × negative control pairs. The query-sample and sample-query GI values for each sgRNA pair were averaged to obtain the sgRNA-level GI. For each gene pair, all sgRNA × sgRNA pairs corresponding to those genes (generally 2×2 except in cases where sgRNAs were filtered) were averaged to obtain the gene-level GI. Negative control gene pairs were simulated by calculating the GI from all combinations of 2 negative control sgRNAs. sgRNA and gene level GIs are provided in Table S4 and S6 respectively.

##### Handling of GIs in which primary phenotypes are positive

When both single perturbations result in a fitness defect, positive and negative GIs can easily be interpreted as buffering or SSL, respectively. When one or both of the perturbations results in increased fitness, however, the interpretation becomes more difficult. Several approaches have been proposed, including changing the sign of the GI value according to the sign of the expected double phenotype (*71*), but these methods necessarily result in discontinuous data and complicate downstream clustering and analysis. For this reason, we did not change the sign of GIs and refer to all positive and negative GIs as buffering and SSL for simplicity.

#### Figure 1

##### *GI map clustering and annotation* (GI_map_all_figures)

For the GI map and Perturb-seq correlation map (Figures 1C, S3, and S4A; see also Figure 3 methods), clustering was performed by average linkage hierarchical clustering using Pearson correlation as the distance metric (SciPy hierarchy package). Principled annotation was performed similar to (*18*) using updated custom Python 2.7 scripts (will be available as github.com/mhorlbeck/hierarchical_annotation.py). Briefly, all DAVID 6.8 default functional annotations ((*40*, *41*); accessed 12/17/2018), including GO BP/MF/CC, BIOCARTA, COG, KEGG, INTERPRO, OMIM, PIR, SMART, and Uniprot, were obtained for each gene in the CRISPRa GI map. Nodes were considered enriched for a given annotation if the log hypergeometric P-value for gene leaves in that node was less than or equal to −7.5 and the node was more enriched for the term than any other node. In cases where a parent of a given node was also enriched for a term and the gene sets overlapped by over 90% or the terms were redundant with a Cohen’s kappa over 0.4, the child node was folded into the parent. Full annotation names were shortened for figures. All node annotations, member genes, and short names are provided in Table S7.

To characterize the relationship between GI and Perturb-seq correlations for specific genes, principal components analysis was performed (scikit-learn 0.20.2). The slope of the relationship was then calculated from the first principal component.

### Perturb-seq analysis and modeling of transcriptional genetic interactions

#### Availability

All analysis of Perturb-seq experiments was performed using a custom package built in Python. This package and the associated notebooks (<u>names appear underlined</u>) used to process the data and generate all the figures will be released alongside the paper.

#### Cell barcode and UMI calling, perturbation calling, data normalization and averaging

(GI_call_barcodes and GI_generate_populations)

Raw sequencing data from 10x experiments was converted to UMI count tables using the default settings of Cellranger 2.1.1. Perturbation identities were called from separate targeted sequencing of Perturb-seq barcodes as described in (*29*). In a departure from the standard 10x workflow, we called perturbation identities for any cell barcode with greater than 2000 UMIs, as Cellranger tended to discard cells bearing perturbations that induced strong growth defects. Any cell barcode passing this UMI threshold and bearing an unambiguous unique perturbation barcode was deemed a cell (i.e., no cell doublets or cells that had received multiple distinct lentiviral integrations) and included in all subsequent analyses.

Normalized expression data for each cell were then derived by (1) equalizing UMI counts across cells, and (2) *z*-normalizing the expression of each gene with respect to the mean and standard deviation observed in control cells bearing nontargeting sgRNAs within the same gemgroup (i.e. same lane on the 10x), as described in (*29*). This normalization does two things: (1) It sets the average expression profile across all genes to be the all-zero vector for unperturbed cells—that is, unperturbed cells define the origin of the resulting coordinate system; and (2) because the normalization is performed within gemgroups, it acts as a simple means of correcting for any batch effects that appear across gemgroups.

For each perturbation (i.e. combination of sgRNAs, representing a distinct genetic background), pseudo-bulk average profiles of normalized and unnormalized UMI count data for all genes were then obtained by averaging across all cells containing that perturbation.

#### Figure 2

##### *Technical* performance of sgRNAs (<u>GI_sgRNA_performance</u>)

Activation performance was computed either as the fold activation (the ratio of the average number of UMIs in perturbed cells vs the average number of UMIs in control cells bearing non-targeting sgRNAs) or the difference (of average UMI counts of the target in perturbed and control cells).

When comparing activation in the A and B position of the vector, cells contain sgRNAs in either of the below configurations:

|  |  |  |
| --- | --- | --- |
| **single_A** | **pos A:** targeting sgRNA | **pos B:** non-targeting control sgRNA |
| **single_B** | **pos A:** non-targeting control sgRNA | **pos B:** targeting sgRNA |

To compare the number of differentially expressed genes as a function of activation level, we executed the following procedure. We restricted our attention to genes with expression greater than 0.25 UMI per cell on average within the experiment. We then compared the distribution of normalized expression within each perturbed cell population to its normalized expression within unperturbed cells using a two-sample Kolmogorov-Smirnov test. *p*-values were corrected for multiple hypothesis testing by the Benjamini-Yekutieli procedure at an FDR of 0.001. The plot in Fig. 2C shows the number of differentially expressed genes called by this procedure as a function of activation level for all perturbations with targeting sgRNAs in the A position and non-targeting control sgRNAs in the second position.

To compare the number of differentially expressed genes for sgRNAs targeting the A and B positions, we (randomly) downsampled each the single_A- and single_B-containing cell populations to contain the same number of cells (whichever number was lower) and called differentially expressed genes as above.

##### *Example gene expression heatmaps* (<u>GI_CBL-CNN1_example</u>)

To identify genes that were differentially expressed across the three genetic backgrounds (single perturbation **a**, single perturbation **b**, and doubly-perturbed cells **ab**), we used a random forest classifier as described in (*72*). Briefly, each cell was used as a training data point, and the classifier was then trained to predict the genetic perturbation a cell contained from its gene expression profile. All genes with mean expression greater than 0.05 UMI per cell were considered as possible distinguishing features. The figures show the top 30 genes with the highest predictive power. Notably, this procedure will tend to identify genes that show large, reproducible differences in expression across condition.

##### *Term enrichment* (<u>GI_CBL-CNN1_example</u>)

Differentially expressed genes in CBL/CNN1 cells were scored with the Kolmogorov-Smirnov test as described above, and then the top 50 genes by *p-*value were submitted to Enrichr (*67*) for term enrichment in the ARCHS4 database of cell-type-specific gene expression (*66*).

##### *Measuring off-target activation in the neighborhood of the targeted gene* (<u>GI_activation_of_neighboring_genes</u>)

To evaluate off-target activation, we used a bootstrap test to evaluate whether the mean expression of genes in the neighborhood of the targeted gene was altered. Specifically, let ***y*** and ***z*** denote the observed vectors of (unnormalized) expression of a given gene in perturbed and unperturbed cells. We wish to test whether *μ*_*y*_ = *μ*_*z*_. To do so, let 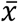 be the mean over all observations and define shifted distributions 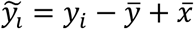 and 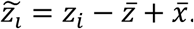 We draw 10,000 bootstrap replicates from 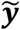 and 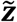 and test for equality of means in each replicate (***y***\*, ***z***\*) via a t-statistic:

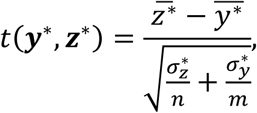

where *n* and *m* are the number of unperturbed and perturbed cells. A *p-*value is assigned as

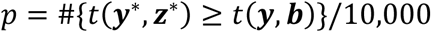

For genes for which we reject the hypothesis at a significance of *p* < 0.05, we then present log_2_ fold changes relative to unperturbed cells. In the figure we mark perturbations that induce large changes in total transcriptome size (<80% of the size in unperturbed cells) with a “#” symbol, as this may indicate widespread changes in gene expression that are not the result of direct sgRNA-induced effects. We applied this testing procedure to the 20 nearest neighbors of the targeted gene in each direction on the chromosome.

#### Figure 3 (GI_optimal_umap)

##### Constructing mean expression profiles

Average normalized gene expression profiles were obtained for all 287 perturbations in the Perturb-seq dataset containing all genes with mean expression level of 0.25 UMI or higher per cell (∼4800 genes). Expression of each gene was then standardized by dividing by the standard deviation across all mean expression profiles. (The average normalized gene expression is close to 0 for all genes by construction since most genes on average are not strongly perturbed away from the unperturbed state.)

##### Clustering of mean expression profiles

Expression profiles for all perturbations were clustered using HDBSCAN with the following parameters: metric=’correlation’, min_cluster_size=4, min_samples=1, cluster_selection_method=’eom’, alpha=1. HDBSCAN only places perturbations into clusters that are “stable”: that is, they are supposed to be somewhat resilient to choices of parameters and subsampling of the data. Profiles that are not part of a stable cluster are left as unclustered outlier points and are gray in the UMAP figure.

##### UMAP projection

UMAP dimensionality reduction was performed on the mean expression profiles using the following parameters: metric=‘correlation’, n_neighbors=10, min_dist=1, spread=2. UMAP is a randomized algorithm, so the precise results depend on the random seed used. Dimensionality reduction was thus performed 10,000 times using different random seeds. For each run, the distance correlation (*73*) was computed between the matrix of mean expression profiles and the reduced matrix of all two-dimensional positions after UMAP dimensionality reduction. We then selected the projection with the highest distance correlation as being most representative of the original high-dimensional data. Points were then colored according to cluster membership determined above.

##### Developmental markers

In the UMAP, expression of different lineage-specific markers were used to define perturbations that were primed towards different differentiation states:

Erythroid: *HBG1*, *HGB2*, *HBZ*, *HGA1*, *HBA2*, *GYPA* (CD235a), *ERMAP*

Granulocyte: *ITGAM* (CD11b), *CSF3R*, *LST1*, *CD33*

The expression of these and other markers are summarized for perturbations in the erythroid and granulocyte clusters in Figure S8F. Note that lowly expressed transcripts and non-protein markers cannot be reliably detected by single-cell RNA seq, so some of the canonical clusters of differentiation (CD) surface markers cannot be measured. For each category, the score was defined as the mean normalized expression of genes in the marker panel.

##### Cell cycle scores

Cell cycle positions were called for each cell using scores derived from panels markers specific to each cell cycle stage as in (*72*). The relative occupancy of cells in each cell cycle stage was then computed for each perturbation. The heatmap of deviations in Figure 3G shows the percentage change in each cell cycle stage relative to the distribution observed in control cells bearing non-targeting sgRNAs. The deviation shown in the paired heatmap shows the correlation distance between occupancy profiles for the given perturbations and unperturbed cells: i.e., the fractions in each cell cycle stage and in each condition are collected into vectors **x** and **y** and the distance is computed as (1 − 𝜌(𝒙, 𝒚))/2.

##### Term enrichment in Figure S8A-E

Perturbations in each of the two clusters were collected together and an average normalized expression profile was computed. Genes with expression greater than 1 in this profile (indicating that their expression was at least one standard deviation above the range observed in unperturbed cells within the cluster on average) were then submitted to Enrichr for term enrichment. The reported scores are Enrichr’s “combined scores” within the indicated databases.

##### Marker gene expression Figure S8F

Specific markers were chosen primarily from (*74*) or (*75*) based on restricted expression within the branches below the common myeloid progenitor (i.e. restricted to either cells derived from proerythroblasts or myeloblasts). A handful of additional markers were included based on strong expression in the data set and restricted expression based on literature searches:

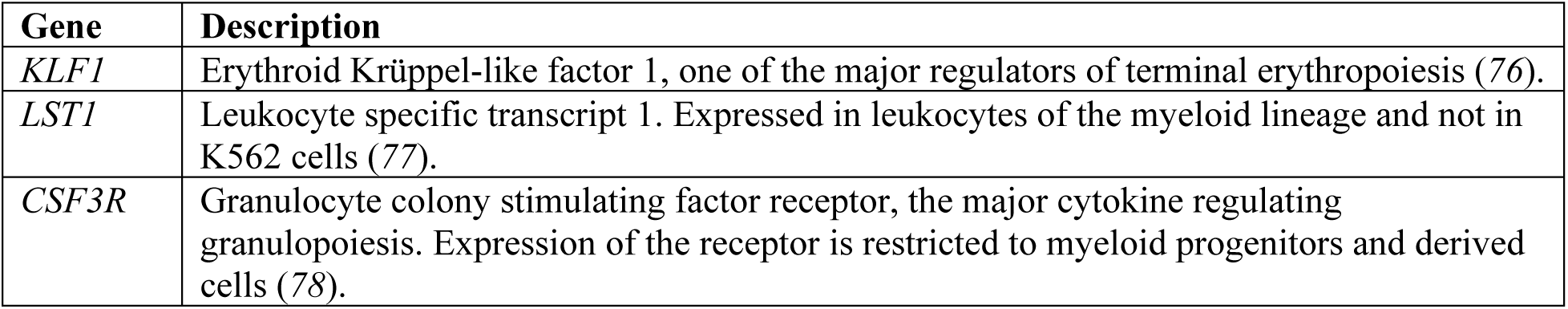

Throughout the text we refer to the clusters of cells expressing these markers as the “erythroid cluster” and the “granulocyte cluster.” This terminology is a shorthand, and the precise placement of these cells in the classical hematopoietic hierarchy is unclear. They may in fact represent transitional, primed, or intermediate states, as indicated by markers such as CD71 (whose expression rises and then falls during erythropoiesis; (*75*)), and CD45/CD123 (used to distinguish erythroid and myeloid progenitors; (*75*)).

To produce the heatmap, the unnormalized average expression profiles for the perturbations were first scaled so that the total UMI count was the same as observed in control cells bearing non-targeting sgRNAs (i.e. we scaled out any global reduction in transcriptome size induced by the perturbation). The heatmap shows log_2_ fold enrichments relative to unperturbed cells. The heatmap is clustered according to the data presented (so the separations of myeloid and erythroid markers and of the two UMAP clusters (*53*) is not forced).

##### Apoptosis scores

Anti-and pro-apoptotic genes were taken from the Qiagen Human Cell Death Pathway Finder RT^2^ Profiler PCR Arrays. Specifically, we used:

Pro-apoptotic: *ABL1, APAF1, ATP6V1G2, BAX, BCL2L11, BIRC2, CASP1, CASP3, CASP6, CASP7, CASP9, CD40, CD40LG, CFLAR, CYLD, DFFA, FAS, FASLG, GADD45A, NOL3, SPATA2, SYCP2, TNF, TNFRSF1A, TNFRSF10A, TP53*

Anti-apoptotic: *AKT1, BCL2, BCL2A1, BCL2L1, BIRC3, CASP2, IGF1R, MCL1, TNFRSF11B, TRAF2, XIAP*

Genes with average expression less than 0.05 UMI per cell were dropped. The total apoptosis score was the mean normalized absolute expression of these genes, representing the total pro- and antiapoptic activity in cells.

##### *Comparisons to erythroid and megakaryocyte developmental time courses* <u>(GI_comparison_to_timecourse</u>)

We compared some of our transcriptional signatures to RNAseq profiles taken from various time points of human CD34+ mobilized peripheral blood cells that had been induced to differentiate through erythropoietin or thrombopoietin treatment. To compare this time course experiment to our Perturb-seq experiment, we first subsetted our transcriptional profiles to genes represented in both data sets with mean expression greater than 0.1 UMI per cell in the Perturb-seq experiment, and then normalized transcriptional profiles from both experiments via the transformation 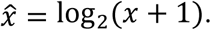 Within the time course data sets, we then characterized genes by their absolute differential expression between first and last time points (i.e. we computed 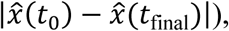 and chose the top 5% of genes by this metric as a signature of differentiation. We then computed Spearman correlations between the time course profiles of these genes at each time point to the perturbation-induced expression profiles in our Perturb-seq experiment. As the null expectation is not necessarily a correlation of 0 (since K562 cells are a hematopoietic cell line and likely share some features with CD34+ cells), so we provide the results for unperturbed K562 cells as a baseline.

#### Figure 4

##### *Fitting the model by robust regression* (GI_model_fits)

To analyze each double perturbation, we consider pseudo-bulk average RNAseq profiles for three different populations of cells defined as below:

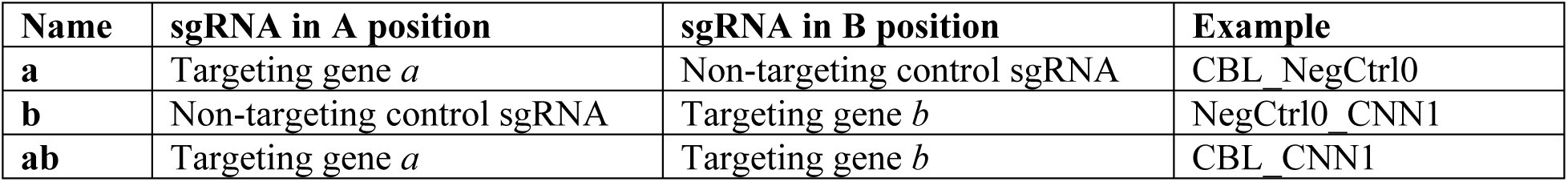

| Name | sgRNA in A position | sgRNA in B position | Example |
| --- | --- | --- | --- |
| **a** | Targeting gene *a* | Non-targeting control sgRNA | CBL_NegCtrl0 |
| **b** | Non-targeting control sgRNA | Targeting gene *b* | NegCtrl0_CNN1 |
| **ab** | Targeting gene *a* | Targeting gene *b* | CBL_CNN1 |

Each genetic perturbation induces changes in expression relative to unperturbed control cells. We can therefore equivalently view, for example, the expression profile **a** in terms of the deviations it induces relative to unperturbed cells:

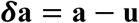

where **u** is the gene expression observed in unperturbed cells. By construction, our normalization procedure (see “Cell barcode and UMI calling, perturbation calling, data normalization and averaging” above) makes this profile all 0 on average for control cells. Thus if we use the pseudo-bulk average normalized expression profiles, unperturbed cells define the origin of the coordinate system and **a** and 𝜹**a** are equivalent.

Our model of genetic interaction then seeks to decompose the perturbation induced by **a** and **b** together in terms of the action of each alone:

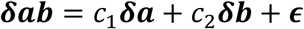

To be clear, here 𝜹**ab**, for example, is the vector of normalized gene expression observed in **ab**- perturbed cells, *c*_1_ and *c*_2_ are constants that are fit to the data, and 𝝐 is a (vector) error term that collects deviations from the overall fit. Geometrically, this can be thought of as trying to best explain the path between unperturbed cells and doubly-perturbed cells as a linear superposition of the two single perturbations.

The goal of fitting the model is to summarize the overall average behavior when perturbations combine, so the coefficients *c*_1_ and *c*_2_ must necessarily average effects over the many genes composing the profiles. Standard least squares regression can be arbitrarily corrupted by outliers (e.g. single genes that undergo massive induction). To reduce this effect, we fit the model coefficients using robust regression, specifically the Theil-Sen estimator (fit on 10,000 random subsamples of 1,000 genes at a time, scikit-learn estimator TheilSenRegressor with parameters fit_intercept=False, max_subpopulation=1e5, max_iter=1000, random_state=1000). We obtained similar results using the implementation of robust linear models in the statsmodels package with the Tukey biweight, but all results in the paper are calculated using the Theil-Sen estimator.

When fitting the model we used all genes with mean expression greater than 0.5 UMI per cell (∼2,800 genes).

##### *Evaluating model fit* (<u>GI_model_fits</u>)

Model fit was evaluated in two ways. The first was the simple *R*^2^ score, which measures:

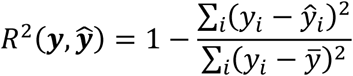

where 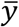 is the sample mean and in this case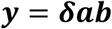 and 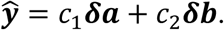

However, because robust regression by definition does not minimize *R*^2^, this is an imperfect metric by which to gauge model fit. In several instances we also evaluated fit via distance correlation (*73*). This metric varies between 0 and 1 and has several advantages over Pearson correlation:

1. it can quantify both linear and nonlinear dependence
2. *dcor*(𝒙, 𝒚) = 0 ⇔ 𝑥 is independent of *y*
3. because it is a function of pairwise distances among rows, *dcor*(𝒙, 𝒚) for example still makes sense when 𝒙 is a vector (i.e. a univariate statistic) and 𝒚 is a matrix (i.e. a multivariate statistic) with the same number of rows (observations) as 𝒙
4. again likely because it is a function of pairwise distances among rows, it in our hands is less sensitive to the effects of individual outlier features (genes)

One disadvantage is that distance correlation is not signed, so positive and negative relationships are not distinguished. In this case, to evaluate fit we computed

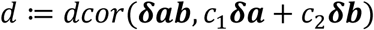

Larger errors will result in smaller values of *d*, which we use to evaluate the degree of neomorphic (i.e. unexpected) behavior. We used the implementation in the dcor Python package.

##### *Example gene expression heatmaps* (<u>GI_DUSP9-MAPK1-ETS2_example</u>)

These were constructed as described above for Figure 2, except that 100 genes are displayed and we only considered genes with mean expression greater than 0.5 UMI (as this was the threshold used for model fitting).

##### *Orienting buffering interactions* (<u>GI_orienting_buffering_interactions</u>)

We looked for interactions that were asymmetrical – i.e. where one single perturbation accounted for substantially more of the double’s behavior than the other. To measure this behavior (examined further in Figure 5), we computed an index of asymmetry as |log_10_(*c*_1_/*c*_2_)|. We considered all interactions where this metric exceeded 1.25 and that had fitness GI scores of 3 or more. In the figure we draw an arrow pointing from the gene with the stronger influence on the double to the one with the weaker influence. The colors were added by taking the expression data as normalized in Figure 3 and clustering it by hierarchical clustering (using the linkage given by setting method=’average’ and metric=’correlation’) to identify perturbations with similar transcriptional profiles.

##### Additive model

In Figure S9, we compared the linear model fit to the additive expectation given by

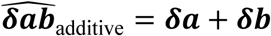

##### *Comparing expression correlation and coefficient magnitudes* (<u>GI_dimensionality_reduction_and_statistics</u>)

Expression data was normalized (as in Figure 3) for all of the “single_A” perturbations (i.e. all perturbations with a targeting sgRNA in the A position of the vector and a non-targeting control sgRNA in the B position). Figure S9C then examines the extent to which correlation of these expression profiles is predictive of interaction as determined by the magnitude of the model coefficients.

##### *Evaluating effects of partner upregulation* (<u>GI_model_fits</u>)

Some interactions (like the ETS2/MAPK1 example) may partly arise when, for example, perturbation of gene *a* also leads to upregulation of gene *b*, so that when both *a* and *b* are perturbed together there is a direct synergy. To evaluate this across our dataset, we compared activation levels within singly- and doubly-perturbed cells. To do this we computed the following ratios:

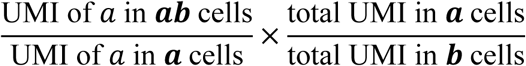

The second factor scales out apparent changes in activation level due to global changes in transcriptome size. The figure plots the log_10_ of these ratios.

#### Figure 5 (GI_dimensionality_reduction_and_statistics)

Figure 5 presents an adaptation of the OneSENSE method (*60*) that uses UMAP (*53*) instead of t-sne as the core dimensionality reduction method. Broadly speaking, the goal is to visualize structure among the interactions in two distinct ways:

(1) in terms of the quantitative model of phenotype we infer in Figure 4
(2) a “model-free” view based on how similar the transcriptional profiles are to each other

The comparison then identifies commonalities and differences between these viewpoints. (E.g. many, but not all, strong synthetic lethal interactions arise from single perturbations that induce similar transcriptional changes.) Accordingly, we constructed transcriptional profiles for each double interaction **ab** and the constituent single interactions **a** and **b** as in Figure 4. We then defined two axes by projecting two sets of variables to a single dimension using UMAP (*53*):

**x-axis:**

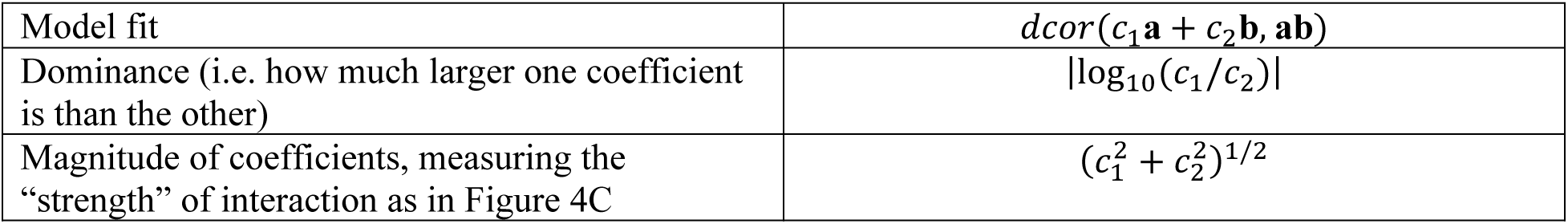

**y-axis:**

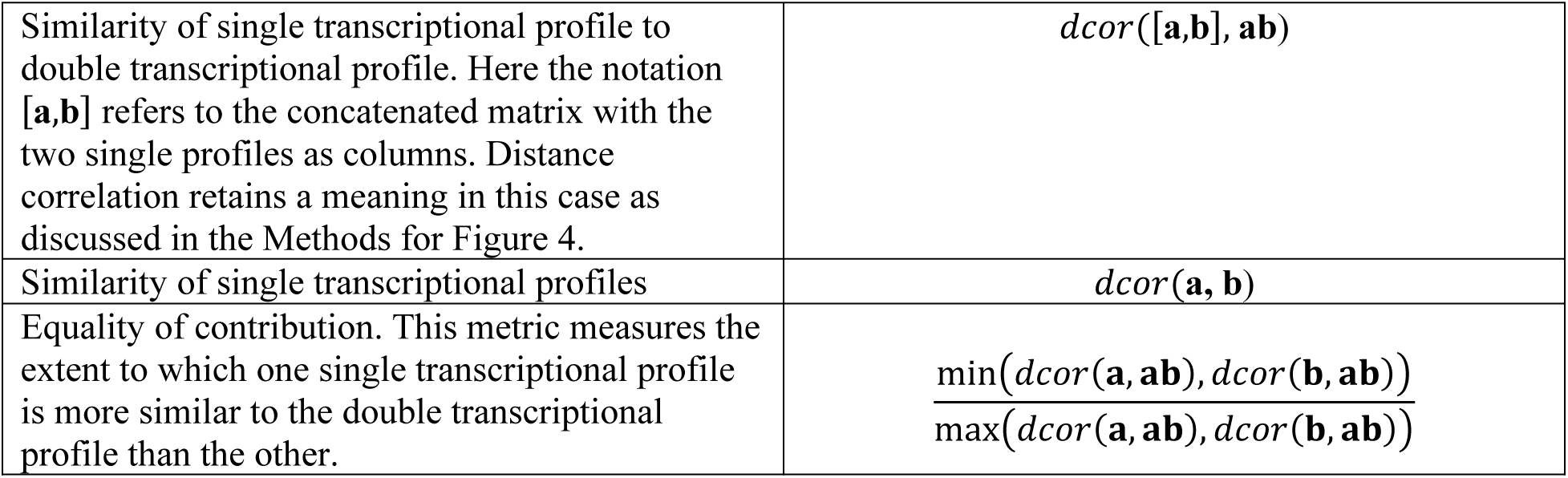

Each of these features was normalized to the same range by dividing by the standard deviation across all of the perturbations considered. The projection was performed using UMAP with parameters: n_neighbors=5, min_dist=0.05, spread=0.5. As the results from UMAP depend on random seed, we chose the specific visualization presented as follows. First, we performed the reduction 10,000 times. In each iterate, we counted the number of clusters formed for the *x* and *y* axes, and qualitatively assessed what number of clusters yielded an appropriate tradeoff between interpretability and granularity (i.e. do not subdivide so much that every interaction is unique, and do not average so much that the clusters were not informative). For each axis we then chose the projection with the chosen number of clusters that had the highest distance correlation with the original three-dimensional data.

#### Figure 6 (GI_DUSP9-MAPK1-ETS2_example)

##### Single-cell UMAP projections

To identify genes that varied across the genetic backgrounds shown, we used the same random forest classifier approach as used in Figure 2 (to make gene expression heatmaps), selecting 200 genes with expression greater than 0.5 UMI per cell. We then projected the matrix of single-cell normalized gene expression data (for all cells from the indicated genetic backgrounds) to two dimensions using UMAP (n_neighbors=10, metric=’euclidean’).

##### Deriving a principal curve and local median filtering

As part of performing dimensionality reduction, UMAP constructs a weighted similarity graph among all cells considered. To identify a principal curve, we reduced this graph to a single dimension using Laplacian eigenmaps, taking this first component as representing the dominant axis of variation in the dataset.

The principal curve allows single cells to be ordered, and hence averaged together. All expression profiles are made by local median filtering: (1) we order the cells along the principal axis (which runs from *t* = −0.0242 to *t* = 0.0273 (2) for each cell along this curve, we compute the median expression over all cells in its neighborhood (distance < 0.001).

The heatmap shows these median filtered data for the genes used in the UMAP projection described above. To distinguish different accumulation patterns, each gene/column (which was itself the *z-*normalized expression data) was further normalized to a 0-1 scale by performing the transformation

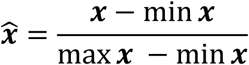

In the panels showing the expression of specific markers, “normalized expression” refers to the *z*-normalized expression described in “Cell barcode and UMI calling, perturbation calling, data normalization and averaging” while “expression accumulation” refers to the 0-1 scaled versions.

### GI prediction analysis

#### Figure 7 (GI_prediction)

##### Predicting genetic interactions

In the analytical approach we have applied to quantify fitness-based genetic interactions (*18*), individual GI scores are not directly observable since they are the result of a fitting procedure that ultimately depends on all of the pairwise fitness measurement. This highlights the general difficulty of predicting GIs: the interactions that are the most interesting are often the ones that are in the tails of the distribution, and are hence the least likely to follow average behavior.

Our procedure for predicting GIs is thus actually a procedure for predicting the distribution of all raw pairwise fitness measurements, which we then use to recompute estimated GI scores. To allow for comparison with past work we have retained the GI estimation procedure from (*18*), but we will highlight an alternative route below as well.

The general workflow of our prediction approach is shown in Figure S11.

(1) We randomly sample fitness measurements for interaction pairs from the full fitness data set at a given sampling rate. For example, at 10% sampling we would sample 10% of pair fitness measurements (we treat **a-b** and **b-a** pairs as identical when doing this), as well as all of the single perturbation fitness measurements.
(2) Fitness GIs ultimately result from double perturbations that are “surprisingly” more or less fit than expected given the two single fitness phenotypes. To define an expected fitness when combining perturbations, we take the fitness measurements from above and fit a quadratic relationship:

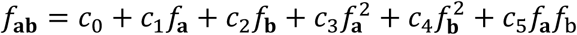

where, for example, 𝑓_**a**_ is the fitness of cells with single perturbation **a**. We enforced symmetry in this fit by adding equivalent “**ba**” pairs for each **ab** interaction, so that *c*_1_ = *c*_2_ and *c*_3_ = *c*_4_. The notable deviation of this fitting procedure from that in (*18*) is that that paper performed the fit on a per gene basis (i.e. each gene had its own fitness expectation). Here we perform randomized measurements over all pairs and so don’t have full fitness profiles for every gene.
(3) We then convert the raw fitness measurements that we have to unexplained “deltas”:

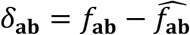 Where 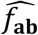 is the expectation from the previous step. These deltas represent deviations from expected fitness, and hence behave similarly as GI scores (as can be seen by comparing the “True deltas” and “True GI map” panels in Figure S11A).
(4) We then predict unobserved deltas using a “recommender system” approach. Many such algorithms have been developed in the context of trying to predict users’ shopping preferences. Analogous to the scenario we consider here, these predictions often have to be made in regimes with very sparse measurements (e.g. a given user has generally purchased only a tiny fraction of available items). The specific approach we use exploits matrix completion, the two-dimensional analog of compressed sensing. Briefly, matrix completion methods take a limited set of measured entries from a matrix and “completes” them to yield an estimate of the whole matrix. Since without constraints this problem is ill-posed, matrix completion approaches will for example assume that the matrix is low rank, which under certain conditions allows it to be perfectly reconstructed even without observing all of the entries. (A low rank matrix has a limited number of degrees of freedom, which implies relationships among the entries that allow them to be perfectly inferred.) A survey of these methods can be found in (*79*). Though matrix completion can be used alone to make predictions from the fitness measurements, methods also exist to leverage side information to enhance predictive power by suggesting how rows and columns might relate to each other. In the shopping context, side information might be for example include a user’s age or geographical location, or a movie’s genre or plot keywords. Given the correlation we observed between transcriptional profile correlation and GI score (fig. S6B), we reasoned that the pairwise correlation among these profiles would serve as useful side information to constrain reconstruction. To start, we generated an expression matrix with all single perturbations (i.e. with targeting sgRNAs in the A position and non-targeting control sgRNAs in the B position) as rows and the unnormalized expression of all genes with mean expression greater than 0.2 UMI per cell as columns. Each gene/column was then normalized to the same scale by performing the transformation

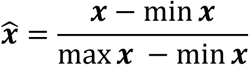 We then computed the pairwise correlation matrix of this expression matrix among all genes Σ. We reasoned that an estimate for the deltas could then be found in the form

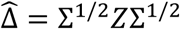

where 𝑍 is an unknown matrix that we fit and Σ^1/2^ is the matrix square root of Σ (unique because the covariance matrix is positive definite). In effect, this posits that the GIs are a “reweighting” of the observed pattern of correlations among transcriptional profiles of genes. As mentioned above, we sought a low-rank 𝑍. Specifically, we sought to solve the following problem:

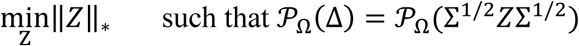

where the 𝒫 condition indicates that the two matrices agree on the set of entries Ω that have been measured directly and ‖𝑍‖_∗_ = ∑𝜌_𝑖_denotes the nuclear norm with 𝜌_𝑖_ the singular values of 𝑍. We solved this problem by applying the Maxide method (*80*) which efficiently solves a relaxed problem:

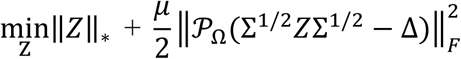

where ‖⋅‖_𝐹_ denotes the Frobenius norm. In all of the results we present we set the Lagrange multiplier *μ* = 0.0005. If 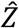 is the result of this optimization, our final estimate is then

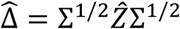
(5) We then convert this estimate back to an estimate for raw fitnesses by adding back the expected fitness for the pair:

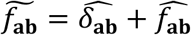
(6) Finally, we estimated GI scores according to the fitting procedure of (*18*), except for a difference in normalization. The procedure presented there normalizes the range of GI scores by the standard deviation in fitness observed when a given sgRNA is paired with a series of negative control sgRNAs (essentially scaling observed the fitness variation due to genetic perturbations by the amount of variation expected to arise due to technical variation). We did not perform this normalization because obtaining these scale factors experimentally would require many more pairwise fitness measurements to be made, which explains why the GI scale seen here is different than in Figure 1. All comparisons in Figure 7 are made to a recomputed version of the GI map made without these normalizations, but with access to the full set of pairwise fitness measurements. It is worth noting that it could also be reasonable to skip this step and the previous one, and merely use the estimate 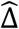 directly as a measure of GIs. In fact, our performance estimating 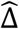 is better, likely simply because it involves fewer steps. Whether it makes sense to fit the expectations for fitness deviation (Step 2) on a per gene basis or to all genes simultaneously may vary depending on context, and we present the results as is for comparison with past work.

##### Evaluating performance

We evaluated performance in three ways. First, to evaluate raw predictive performance over all interactions, we measured the Spearman correlation (testing similarity of rank order) between predicted GIs and true GIs.

Second, we tested similarity at the level of GI profiles for each gene. This we evaluated via cophenetic correlation: i.e. we computed the (Pearson) correlation between vectors containing all pairwise distances of columns in the predicted and true GI maps. Performance varied by distance metric—in Figure 7 we present the results when columns are compared using cosine distance, which has been used to cluster GI maps in the past (*18*). In Figure S11 we compared columns using Euclidean distance, which is somewhat better preserved.

Finally, we wished to evaluate the ability to predict strong GIs. We did this by testing recall. We defined GIs as strong if they fell either below the 5^th^ percentile or above the 95^th^ percentile of all GI scores. The recall is the fraction of these that then fell in the same bins in the predicted GI map. Random sampling will discover these with uniform probability, so e.g. 5% random sampling will be expected to find 5% of strong GIs.

Matrix completion can also be performed without side information. To assess the improvement in predictive power made by using the Perturb-seq side information, Figures S11E-G show comparisons of the data from Figure 7D-F to predictions made with the side information set to the identity matrix.

##### Routes to improving performance

In principle performance could be improved using biased sampling or different side information. To illustrate this qualitative point, we employed two simple modifications to the procedure described above.

First, we did non-random sampling of interaction pairs. Following the same reasoning as when constructing our side information matrix above, we thought that a biased sampling of pairs based on the correlation of their profiles might improve performance. We first generated a matrix 𝑀 consisting of all the pairwise distance correlations among transcriptional profiles (which were constructed as in Step 4 above). We then centered this matrix by subtracting the median distance correlation over all pairs. Finally, we performed random sampling of its entries according to their magnitude as in the LELA (Leveraged Element Low-rank Approximation) algorithm in (*81*). Briefly, this approach approximates the leverage scores of each entry in the matrix (which in a sense measure how much information a given entry carries about the matrix’s structure) to construct a biased sampling distribution. Practically speaking, this will less frequently sample regions of the matrix that behave similarly (like the coherent blocks seen in GI maps or in pairwise correlation matrices).

Second, we constructed a different side information matrix. This matrix Σ_2_ was constructed in the same was as the original one, but with the z-normalized gene expression data instead of the unnormalized UMI counts and with genes with standard deviation greater than 6 across perturbations filtered out (to remove genes showing extreme perturbation-specific induction). As the final side information input, we used Σ^1/2^ + Σ_2_^1/2^. The minimizer 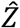 is then forced to be consistent with two different scalings of our gene expression data—one “raw” scale, and one that has been normalized so that it expresses deviations from unperturbed cells.

Each of these changes individually improves performance, and the combination, which we present in Figure S11D, improves performance further. Though the gains we observe are modest and the reasoning behind each of these alterations is largely ad hoc, they do show qualitatively that improvements in performance are possible through rational interventions. With larger data sets we envision that the principles needed to design both better sampling strategies and more informative side information can be learned.

##### *Downsampling analysis* (<u>GI_downsampling_analysis</u>)

Downsampling analyses were performed by randomly the given number of cells from each genetic perturbation background 50 times, and then re-executing the identical clustering analysis used in Figure 3 (“Clustering of mean expression profiles”) and the model fitting procedure from Figure 4.

To evaluate the similarity of measured transcriptional phenotypes, we computed the cophenetic correlation (Pearson correlation of all pairwise distances computed using the ‘correlation’ metric) between each downsampled population and all cells we gathered.

To evaluate robustness of the model fitting procedure, we compared the measured coefficient magnitude (measuring interaction strength) and dominance (measuring asymmetry of contributions of the two single perturbations to the double’s phenotype). As genes with low expression are more sensitive to noise when fewer cells are measured, we also refit model coefficients using only genes with mean expression greater than 1 UMI per cell.

## References

1. J.-M. Claverie, What If There Are Only 30,000 Human Genes? Science. 291, 1255–1257 (2001).

2. M. Costanzo et al., The genetic landscape of a cell. Science. 327, 425–431 (2010).

3. M. Costanzo et al., A global genetic interaction network maps a wiring diagram of cellular function. Science. 353 (2016), doi:10.1126/science.aaf1420.

4. X. Pan et al., A robust toolkit for functional profiling of the yeast genome. Mol. Cell. 16, 487–496 (2004).

5. K. Takahashi, S. Yamanaka, Induction of pluripotent stem cells from mouse embryonic and adult fibroblast cultures by defined factors. Cell. 126, 663–676 (2006).

6. A. H. Tong et al., Systematic genetic analysis with ordered arrays of yeast deletion mutants. Science. 294, 2364–2368 (2001).

7. A. H. Y. Tong et al., Global mapping of the yeast genetic interaction network. Science. 303, 808–813 (2004).

8. P. Novick, B. C. Osmond, D. Botstein, Suppressors of yeast actin mutations. Genetics. 121, 659–674 (1989).

9. T. Stearns, D. Botstein, Unlinked noncomplementation: isolation of new conditional-lethal mutations in each of the tubulin genes of Saccharomyces cerevisiae. Genetics. 119, 249–260 (1988).

10. C. A. Kaiser, R. Schekman, Distinct sets of SEC genes govern transport vesicle formation and fusion early in the secretory pathway. Cell. 61, 723–733 (1990).

11. S. R. Collins et al., Functional dissection of protein complexes involved in yeast chromosome biology using a genetic interaction map. Nature. 446, 806–810 (2007).

12. M. C. Jonikas et al., Comprehensive characterization of genes required for protein folding in the endoplasmic reticulum. Science. 323, 1693–1697 (2009).

13. X. Pan et al., A DNA integrity network in the yeast Saccharomyces cerevisiae. Cell. 124, 1069–1081 (2006).

14. M. Schuldiner et al., Exploration of the function and organization of the yeast early secretory pathway through an epistatic miniarray profile. Cell. 123, 507–519 (2005).

15. D. Segrè, A. Deluna, G. M. Church, R. Kishony, Modular epistasis in yeast metabolism. Nat. Genet. 37, 77–83 (2005).

16. S. Bandyopadhyay et al., Rewiring of genetic networks in response to DNA damage. Science. 330, 1385–1389 (2010).

17. M. Costanzo et al., Global Genetic Networks and the Genotype-to-Phenotype Relationship. Cell. 177, 85–100 (2019).

18. M. A. Horlbeck et al., Mapping the Genetic Landscape of Human Cells. Cell. 174, 953–967.e22 (2018).

19. M. Boettcher et al., Dual gene activation and knockout screen reveals directional dependencies in genetic networks. Nat. Biotechnol. 36, 170–178 (2018).

20. M. C. Bassik et al., A systematic mammalian genetic interaction map reveals pathways underlying ricin susceptibility. Cell. 152, 909–922 (2013).

21. D. Du et al., Genetic interaction mapping in mammalian cells using CRISPR interference. Nat. Methods. 14, 577–580 (2017).

22. K. Han et al., Synergistic drug combinations for cancer identified in a CRISPR screen for pairwise genetic interactions. Nat. Biotechnol. 35, 463–474 (2017).

23. A. Roguev et al., Quantitative genetic-interaction mapping in mammalian cells. Nat. Methods. 10, 432–437 (2013).

24. J. Rosenbluh et al., Genetic and Proteomic Interrogation of Lower Confidence Candidate Genes Reveals Signaling Networks in β-Catenin-Active Cancers. Cell Syst. 3, 302–316.e4 (2016).

25. J. P. Shen et al., Combinatorial CRISPR-Cas9 screens for de novo mapping of genetic interactions. Nat. Methods. 14, 573–576 (2017).

26. A. S. L. Wong et al., Multiplexed barcoded CRISPR-Cas9 screening enabled by CombiGEM. Proc. Natl. Acad. Sci. U.S.A. 113, 2544–2549 (2016).

27. Y. Liu et al., CRISPR Activation Screens Systematically Identify Factors that Drive Neuronal Fate and Reprogramming. Cell Stem Cell. 23, 758–771.e8 (2018).

28. U. Parekh et al., Mapping Cellular Reprogramming via Pooled Overexpression Screens with Paired Fitness and Single-Cell RNA-Sequencing Readout. Cell Syst. 7, 548–555.e8 (2018).

29. B. Adamson et al., A Multiplexed Single-Cell CRISPR Screening Platform Enables Systematic Dissection of the Unfolded Protein Response. Cell. 167, 1867–1882.e21 (2016).

30. A. Dixit et al., Perturb-seq: Dissecting molecular circuits with scalable single cell RNA profiling of pooled genetic screens. Cell. 167, 1853–1866.e17 (2016).

31. D. A. Jaitin et al., Dissecting Immune Circuits by Linking CRISPR-Pooled Screens with Single-Cell RNA-Seq. Cell. 167, 1883–1896.e15 (2016).

32. P. Datlinger et al., Pooled CRISPR screening with single-cell transcriptome readout. Nat. Methods. 14, 297–301 (2017).

33. J. M. Replogle et al., Direct capture of CRISPR guides enables scalable, multiplexed, and multi-omic Perturb-seq. bioRxiv, 503367 (2018).

34. M. Gasperini et al., A Genome-wide Framework for Mapping Gene Regulation via Cellular Genetic Screens. Cell. 176, 377–390.e19 (2019).

35. L. A. Gilbert et al., Genome-Scale CRISPR-Mediated Control of Gene Repression and Activation. Cell. 159, 647–661 (2014).

36. J.-Y. Youn et al., Functional Analysis of Kinases and Transcription Factors in Saccharomyces cerevisiae Using an Integrated Overexpression Library. G3 (Bethesda). 7, 911–921 (2017).

37. L. M. Sack et al., Profound Tissue Specificity in Proliferation Control Underlies Cancer Drivers and Aneuploidy Patterns. Cell. 173, 499–514.e23 (2018).

38. S. Duffy et al., Overexpression screens identify conserved dosage chromosome instability genes in yeast and human cancer. Proc. Natl. Acad. Sci. U.S.A. 113, 9967–9976 (2016).

39. M. E. Tanenbaum, L. A. Gilbert, L. S. Qi, J. S. Weissman, R. D. Vale, A protein-tagging system for signal amplification in gene expression and fluorescence imaging. Cell. 159, 635–646 (2014).

40. D. W. Huang, B. T. Sherman, R. A. Lempicki, Systematic and integrative analysis of large gene lists using DAVID bioinformatics resources. Nat Protoc. 4, 44–57 (2009).

41. D. W. Huang, B. T. Sherman, R. A. Lempicki, Bioinformatics enrichment tools: paths toward the comprehensive functional analysis of large gene lists. Nucleic Acids Res. 37, 1–13 (2009).

42. M. Ohta et al., Direct interaction of Plk4 with STIL ensures formation of a single procentriole per parental centriole. Nat Commun. 5, 5267 (2014).

43. A. Dongre, R. A. Weinberg, New insights into the mechanisms of epithelial-mesenchymal transition and implications for cancer. Nat. Rev. Mol. Cell Biol. 20, 69–84 (2019).

44. J. Kale, E. J. Osterlund, D. W. Andrews, BCL-2 family proteins: changing partners in the dance towards death. Cell Death Differ. 25, 65–80 (2018).

45. S. Konermann et al., Genome-scale transcriptional activation by an engineered CRISPR-Cas9 complex. Nature. 517, 583–588 (2015).

46. C. A. Joazeiro et al., The tyrosine kinase negative regulator c-Cbl as a RING-type, E2-dependent ubiquitin-protein ligase. Science. 286, 309–312 (1999).

47. S. J. Winder, M. P. Walsh, Calponin: thin filament-linked regulation of smooth muscle contraction. Cell. Signal. 5, 677–686 (1993).

48. GTEx Consortium et al., Genetic effects on gene expression across human tissues. Nature. 550, 204–213 (2017).

49. FANTOM Consortium and the RIKEN PMI and CLST (DGT) et al., A promoter-level mammalian expression atlas. Nature. 507, 462–470 (2014).

50. B. J. Druker et al., Effects of a selective inhibitor of the Abl tyrosine kinase on the growth of Bcr-Abl positive cells. Nat. Med. 2, 561–566 (1996).

51. B. Bruecher-Encke, J. D. Griffin, B. G. Neel, U. Lorenz, Role of the tyrosine phosphatase SHP-1 in K562 cell differentiation. Leukemia. 15, 1424–1432 (2001).

52. R. Kurita et al., Establishment of immortalized human erythroid progenitor cell lines able to produce enucleated red blood cells. PLoS ONE. 8, e59890 (2013).

53. L. McInnes, J. Healy, J. Melville, UMAP: Uniform Manifold Approximation and Projection for Dimension Reduction (2018) (available at https://arxiv.org/abs/1802.03426v2).

54. K. Weiskopf et al., Myeloid cell origins, differentiation, and clinical implications. Microbiol Spectr. 4 (2016), doi:10.1128/microbiolspec.MCHD-0031-2016.

55. E. M. Rabellino, R. L. Nachman, N. Williams, R. J. Winchester, G. D. Ross, Human megakaryocytes. I. Characterization of the membrane and cytoplasmic components of isolated marrow megakaryocytes. J. Exp. Med. 149, 1273–1287 (1979).

56. G. Vinci et al., Immunological study of in vitro maturation of human megakaryocytes. Br. J. Haematol. 56, 589–605 (1984).

57. M. T. Mitjavila-Garcia et al., Expression of CD41 on hematopoietic progenitors derived from embryonic hematopoietic cells. Development. 129, 2003–2013 (2002).

58. M. E. Tanenbaum et al., A complex of Kif18b and MCAK promotes microtubule depolymerization and is negatively regulated by Aurora kinases. Curr. Biol. 21, 1356–1365 (2011).

59. C. J. Caunt, S. M. Keyse, Dual-specificity MAP kinase phosphatases (MKPs). FEBS J. 280, 489–504 (2013).

60. Y. Cheng, M. T. Wong, L. van der Maaten, E. W. Newell, Categorical Analysis of Human T Cell Heterogeneity with One-Dimensional Soli-Expression by Nonlinear Stochastic Embedding. J. Immunol. 196, 924–932 (2016).

61. C. Trapnell et al., The dynamics and regulators of cell fate decisions are revealed by pseudotemporal ordering of single cells. Nat. Biotechnol. 32, 381–386 (2014).

62. H.-T. Cheng et al., in Proceedings of the 1st Workshop on Deep Learning for Recommender Systems - DLRS 2016 (ACM Press, Boston, MA, USA, 2016; http://dl.acm.org/citation.cfm?doid=2988450.2988454), pp. 7–10.

63. D. Feldman et al., Pooled optical screens in human cells. bioRxiv, 383943 (2018).

64. E. Romanov, M. Gavish, Near-optimal matrix recovery from random linear measurements. Proc. Natl. Acad. Sci. U.S.A. 115, 7200–7205 (2018).

65. D. L. Donoho, A. Maleki, A. Montanari, Message-passing algorithms for compressed sensing. Proc. Natl. Acad. Sci. U.S.A. 106, 18914–18919 (2009).

66. A. Lachmann et al., Massive mining of publicly available RNA-seq data from human and mouse. Nat Commun. 9, 1366 (2018).

67. M. V. Kuleshov et al., Enrichr: a comprehensive gene set enrichment analysis web server 2016 update. Nucleic Acids Res. 44, W90–W97 (2016).

68. B. Chen et al., Dynamic imaging of genomic loci in living human cells by an optimized CRISPR/Cas system. Cell. 155, 1479–1491 (2013).

69. K. Weber, U. Bartsch, C. Stocking, B. Fehse, A multicolor panel of novel lentiviral “gene ontology” (LeGO) vectors for functional gene analysis. Mol. Ther. 16, 698–706 (2008).

70. M. A. Horlbeck et al., Compact and highly active next-generation libraries for CRISPR-mediated gene repression and activation. Elife. 5 (2016), doi:10.7554/eLife.19760.

71. M. Kampmann, M. C. Bassik, J. S. Weissman, Integrated platform for genome-wide screening and construction of high-density genetic interaction maps in mammalian cells. Proc. Natl. Acad. Sci. U.S.A. 110, E2317–2326 (2013).

72. B. Adamson et al., A Multiplexed Single-Cell CRISPR Screening Platform Enables Systematic Dissection of the Unfolded Protein Response. Cell. 167, 1867–1882.e21 (2016).

73. G. J. Székely, M. L. Rizzo, N. K. Bakirov, Measuring and testing dependence by correlation of distances. Ann. Statist. 35, 2769–2794 (2007).

74. BD Bioscience CD Marker Handbook, cd_marker_handbook.pdf, (available at https://www.bdbiosciences.com/documents/cd_marker_handbook.pdf).

75. A. Attar, Changes in the Cell Surface Markers During Normal Hematopoiesis: A Guide to Cell Isolation. Global Journal of Hematology and Blood Transfusion (2014), doi:10.15379/2408-9877.2014.01.01.4.

76. M. Siatecka, J. J. Bieker, The multifunctional role of EKLF/KLF1 during erythropoiesis. Blood. 118, 2044–2054 (2011).

77. P. Draber et al., LST1/A is a myeloid leukocyte-specific transmembrane adaptor protein recruiting protein tyrosine phosphatases SHP-1 and SHP-2 to the plasma membrane. J. Biol. Chem. 287, 22812–22821 (2012).

78. K. Tsuji, Y. Ebihara, Expression of G-CSF receptor on myeloid progenitors. Leuk. Lymphoma. 42, 1351–1357 (2001).

79. M. A. Davenport, J. Romberg, An overview of low-rank matrix recovery from incomplete observations. IEEE Journal of Selected Topics in Signal Processing. 10, 608–622 (2016).

80. M. Xu, R. Jin, Z.-H. Zhou, Speedup Matrix Completion with Side Information: Application to Multi-Label Learning, 9.

81. S. Bhojanapalli, P. Jain, S. Sanghavi, in Proceedings of the Twenty-Sixth Annual ACM-SIAM Symposium on Discrete Algorithms (Society for Industrial and Applied Mathematics, 2015; https://epubs.siam.org/doi/10.1137/1.9781611973730.62), pp. 902–920.

